# Histological and gene-expression analyses of pyloric sphincter formation during stomach metamorphosis in *Xenopus laevis*

**DOI:** 10.1101/2024.04.29.591326

**Authors:** Kei Nagura, Takafumi Ikeda, Takashi Hasebe, Yumeko Satou-Kobayashi, Sumio Udagawa, Shuji Shigenobu, Atsuko Ishizuya-Oka, Masanori Taira

**Affiliations:** Department of Biological Sciences, Graduate School of Science, The University of Tokyo, 7-3-1 Hongo, Bunkyo-ku, Tokyo 113-0033, Japan; Institute for Protein Dynamics, Kyoto Sangyo University, Kyoto 603-8555, Japan (present address); Faculty of Life Sciences, Kyoto Sangyo University, Kyoto 603-8555, Japan (present address); Department of Biology, Nippon Medical School, Kyonan-cho, Musashino, Tokyo 180-0023, Japan; Advanced Comprehensive Research Organization, Teikyo University, 2-11-1 Kaga, Itabashi-ku, Tokyo 173-0003, Japan (present address); Misaki Marine Biological Station, Graduate School of Science and Center for Marine Biology, The University of Tokyo, 1024 Koajiro Misaki, Miura, Kanagawa 238-0225, Japan; Tateyama Marine Laboratory, Marine and Coastal Research Center, Ochanomizu University, Kou-yatsu 11, Tateyama, Chiba, 294-0301, Japan (present address); National Institute for Basic Biology (NIBB), Nishigonaka 38, Myodaiji, Okazaki 444-8585, Japan; Department of Biological Sciences, Faculty of Science and Engineering, Chuo University, 1-13-27 Kasuga, Bunkyo-ku, Tokyo 112-8551, Japan (present address)

**Author notes:** Corresponding author: Masanori Taira. These authors contributed equally to this work (Kei Nagura, Takafumi Ikeda).

**Keywords:** *Xenopus laevis*, metamorphosis, remodeling, stomach, pylorus, pyloric sphincter, gene expression, RNA-seq analysis

## Abstract

During anuran metamorphosis from herbivorous tadpoles to carnivorous frogs, the gastrointestinal (GI) tract undergoes drastic remodeling, such as the formation of the stomach-intestine boundary and the development of the pyloric sphincter at the posterior end of the stomach. However, the morphogenetic process and molecular mechanisms of how the pyloric sphincter is formed during metamorphosis, instead of during embryogenesis as in amniotes, are largely uninvestigated. Using the African clawed frog *Xenopus laevis*, we histologically examined the development of the pylorus region from embryonic to froglet stages and performed spatiotemporal gene expression analyses. We found that the pyloric sphincter is formed at a flexure within the pyloric region during metamorphic climax, and that the pyloric and duodenal epithelia, which are morphologically indistinguishable before sphincter formation, become clearly demarcated by the sphincter at the end of metamorphosis. Consistent with these morphological changes, expression domains of a stomach marker *barx1* and an intestine marker *cdx2* overlapped until late metamorphic climax, but became separated after metamorphosis. Despite the absence of the sphincter before metamorphosis, various genes crucial for sphincter formation in amniotes were already expressed in the pylorus region of *Xenopus* embryos. RNA-sequencing analysis at pre-metamorphic and metamorphic-climax stages suggest unappreciated roles of genes, such as those for retinoic acid signaling and various transcription factors, in suppressing or promoting sphincter formation. These data provide histological and molecular insights into the heterochrony of the pyloric sphincter formation in amniotes and anurans.

## Introduction

The morphology of the gastrointestinal (GI) tract in vertebrates is greatly diversified according to feeding habits (Stevens and Hume, 1995). In most anurans, herbivorous larvae (tadpoles) and carnivorous adults (frogs) have morphologically and functionally distinct GI tracts. This is achieved by the remodeling of the larval GI tract, which occurs during thyroid hormone (TH)-induced metamorphosis, such as the shortening of the intestine (Brown and Cai, 2007), the onset of pepsinogen secretion from the stomach (Ishizuya-Oka et al., 1998), and the formation of the pyloric sphincter at the posterior end of the stomach (Smith et al., 2000a). Formation of the pyloric sphincter is crucial for establishing a carnivorous digestive system because it enables functional division of the stomach and intestine to protect the intestine from the acidic gastric juice secreted in the stomach (Chandranath et al., 2014). While morphological and molecular analyses on intestinal remodeling have progressed (Fu et al., 2019; Ishizuya-Oka, 2017; Schreiber et al., 2005; Sterling et al., 2012), the remodeling of the stomach during metamorphosis, including pyloric sphincter formation, is not fully understood.

The vertebrate stomach is generally composed of the anterior (proximal) and posterior (distal) stomach. The anterior stomach is further divided into the cardia, fundus and stomach body, and the posterior stomach is divided into the antrum and the pylorus region, followed by the small intestine comprising the duodenum and ileum. Although these morphological terms are defined according to the adult morphology of humans, we refer to the corresponding regions of the *Xenopus* GI tract with the same terms, following Xenbase Anatomy Ontology (https://www.xenbase.org/entry/anatomy/xao.do?method=display). In amniotes, regionalization of the GI tract along the antero-posterior axis occurs through epithelial-mesenchymal interaction, which induces region-specific expression of transcription factors such as *barx1* in the stomach, *nkx2-5* and *sox9* in the pylorus region, and *cdx2* in the intestine (Le Guen et al., 2015; Smith et al., 2000a). In mice, *Barx1* and *Cdx2* are expressed in the stomach mesenchyme and intestinal epithelium, respectively (Beck et al., 2000; Gao et al., 2009; Kim et al., 2005; Li et al., 2009; Tissier-Seta et al., 1995), making them molecular markers for the stomach and intestine.

Pyloric sphincter formation is characterized by the thickening of the circular muscle layer and the narrowing of the lumen in the pylorus region, followed by the formation of the boundary in the epithelium of the stomach and duodenum (Li et al., 2009; Nieuwkoop and Faber, 1967; Self et al., 2009). In mice and chickens, *Nkx2-5*, *Sox9,* and *Gremlin* are specifically expressed in the pyloric mesoderm and involved in the formation of the pyloric epithelium (Li et al., 2014; Moniot et al., 2004; Smith et al., 2000a; Theodosiou and Tabin, 2005; Udager et al., 2010). Expression of *Nkx2-5* and *Sox9* is induced by the BMP signal, which is downstream of the Shh signal (Roberts et al., 1995; Roberts et al., 1998; Theodosiou and Tabin, 2005). In mice, *Bapx1* (a synonym of *Nkx3-2*) is expressed in the antrum and pylorus region, whereas *Gata3* is specifically expressed in the pylorus region, both of which are required for pyloric sphincter formation (Prakash et al., 2014; Verzi et al., 2009).

In contrast to amniotes, regionalization of the stomach and pyloric sphincter formation during anuran metamorphosis has not been well investigated. *Xenopus laevis*, and more recently *X. tropicalis*, have been used as model anuran species for histological studies of GI-tract morphogenesis (Ikuzawa et al., 2004; Ishizuya-Oka et al., 2003; Nieuwkoop and Faber, 1967; Oinuma et al., 1992; Smith et al., 2000a). As in amniotes, the stomach and duodenum in *Xenopus* are regionalized morphologically and molecularly during embryogenesis (Smith et al., 2000a). However, unlike amniotes, herbivorous *Xenopus* and *Rana* tadpoles have a long posterior stomach with a continuous simple columnar epithelium from the posterior stomach to the duodenum and do not have the pyloric sphincter before metamorphosis (Smith et al., 2000a) (Barrington, 1946). Based on this morphological feature together with the presence of ciliated epithelium and mucosecretory cells, both of which are absent in the duodenum, the posterior stomach of tadpoles is also called the transitional zone (Barrington, 1946; Bodegas et al., 1997; Chalmers et al., 2000). Metamorphosis in *Xenopus* proceeds through pre-metamorphosis (stages 52-55), pro-metamorphosis (stages 56-58), and metamorphic climax (stages 59-65), during which the GI tract and other organs are dramatically remodeled (Brown and Cai, 2007). Although Nieuwkoop and Faber (1967) (Nieuwkoop and Faber, 1967) described that the metamorphosis of the pylorus region begins in the middle of metamorphic climax (stage 61) and completes at stage 64, it is unclear when the posterior stomach and duodenum are regionalized and how the pyloric sphincter is formed during metamorphosis.

Besides the lack of histological descriptions, the molecular mechanism of pylorus formation and regionalization of the stomach and duodenum are poorly understood in anurans. In *Xenopus*, *sox2* expression in the stomach and *Xcad1* and *Xcad2* (synonyms of *cdx2* and *cdx1*, respectively) expression in the intestine are reported at stages 41 (embryonic) and 45/46 (tadpole) (Chalmers et al., 2000). At stage 42, *nkx2-5* and *nkx2-3* are expressed in the pyloric and intestinal mesoderm, respectively, but overlap in the posterior stomach (Smith et al., 2000a), which may be distinct from amniotes (Udager et al., 2010). However, little is known about whether and how other related genes such as *gata3*, *bmp4* and *gremlin* (*grem1*) are differentially expressed in embryogenesis and metamorphosis. Of note, treatment with an antagonist of retinoic acid (RA) can transform the herbivorous stomach of the *Xenopus* larva to an enlarged, carnivore-like one (Bloom et al., 2013). This raises the possibility that suppression of RA signaling plays a pivotal role in remodeling the stomach from herbivorous into carnivorous, but expression patterns of RA-related genes during *Xenopus* metamorphosis remain elusive.

To address the mechanism underlying the morphogenetic process of the pylorus region during anuran metamorphosis, we first performed detailed histological analyses using *Xenopus laevis*. We then conducted expression analyses of genes referred here to as “pylorus/sphincter-related genes,” which include stomach and intestine marker genes, *barx1* and *cdx2*, as well as genes that are involved in the pyloric sphincter formation in amniotes, *nkx2-3, nkx2-5, nkx3-2, gata3*, *sox9, bmp4,* and *grem1,* as mentioned above. Furthermore, to comprehensively search for genes involved in the metamorphosis of the pylorus region, we performed RNA-sequencing (RNA-seq) with regionally dissected GI tracts at pre-metamorphic and metamorphic-climax stages. Based on these analyses, we discuss how the morphogenetic mechanisms of the pylorus region are diversified in amniotes and anurans.

## Materials and methods

### Animals

All experiments using *Xenopus laevis* were approved by the Office for Life Science Research Ethics and Safety, the University of Tokyo (Approval No. 13-06). Adult male and female frogs of *Xenopus laevis* and stage 53 - 60 tadpoles of the J-strain (an inbred line) were purchased from Watanabe Zoshoku (Hyogo, Japan). *X. laevis* embryos were obtained using artificial fertilization and reared until desired stages. Tadpoles, froglets, and adult frogs were used for histological sections and in situ hybridization. The J-strain was used for RT-PCR and RNA-seq.

### Stages

Developmental stages of *Xenopus laevis* were determined according to Nieuwkoop and Faber (1967). Xenbase (http://wiki.xenbase.org/xenwiki/index.php/Xenopus_development_stages) was also used to determine the embryonic/tailbud stages (by stage 44) and tadpole stages (from stage 45 onward). Tailbud embryos at stage 42/43, tadpoles at stage 46/47, pre-metamorphic tadpoles at stages 51 to 53, metamorphic tadpoles at stage 60 to 63, and froglet at stage 66 were used.

### Histological analyses

Histological analysis was performed using the standard method. Feeding of the larvae was stopped one day before fixation. GI tracts were isolated using forceps and a razor blade under a stereo microscope, and contents of GI tracts were removed after fixation, if necessary. Isolated GI tracts were fixed for one day with Bouin’s fixative and stored in 70% ethanol until use. Fixed GI tracts were manually trimmed using a razor blade and the desired regions were embedded in paraffin. Serial sections were cut at 5 µm, stained with hematoxylin and eosin, and observed using a ZEISS Axioplan2 microscope.

### cDNA cloning and synthesis of RNA probes

Either of the *X. laevis* L and or S homeologs with a higher expression level than the other at embryonic stages based on the public RNA-seq datasets (Session et al., 2016) was subjected to cDNA cloning. Coding sequences of *barx1.L, cdx2.L, nkx2-5.S, gata3.S, grem1.L, nkx3-2.L* and *acta2.L* were PCR-amplified from *X. laevis* cDNA libraries or cDNA mixtures, and were inserted into the pCSf107mT vector (Mii and Taira, 2009). pCSf107-BMP4.S-T (a gift from Takayoshi Yamamoto) and pGEMT-Xsox9 (Spokony et al., 2002) were used as *bmp4.S* and *sox9.S* clones. Digoxigenin (DIG)-labeled antisense and sense RNA probes for whole-mount in situ hybridization (WISH) were transcribed from these plasmids using T7 or SP6 polymerases (Roche).

### Whole-mount in situ hybridization

Whole-mount in situ hybridization (WISH) was performed using a standard protocol (Harland, 1991). Embryonic and tadpole GI tracts were isolated as described above, fixed with MEMFA fixative, and hybridized with DIG-labelled RNA probes. BM purple (Roche) was used for chromogenic reaction.

### Section in situ hybridization

Section in situ hybridization (ISH) was performed basically as reported by Hasebe et al. (2006). Isolated GI tracts were fixed with MEMFA fixative for two hours at room temperature, embedded in OCT (optimal cutting temperature) compound (Sakura Finetek), and cryosectioned at 7 µm. Hybridization was conducted overnight at 60 °C. Chromogenic reaction was carried out for one to six days at 23 °C. NBT/BCIP solution (Roche) was used as the chromogenic substrate.

### RNA extraction

The anterior constriction of the stomach body was used as a positional marker to dissect pieces every one millimeter from stages-53 and -60 GI tracts (see Fig. 7A). Pieces from the anterior to posterior positions were serially numbered with prefixes T (tadpole stage) and M (metamorphic stage). For metamorphic and froglet stages (stages 62 and 66, respectively), the flexure and regions anterior and posterior to it was divided into regions 1 (antrum), 2 (pylorus), and 3 (duodenum) (see Fig. 7A). Total RNA was extracted from dissected pieces from three to seven individuals using ISOGEN or ISOGEN-LS (Nippon Gene). Extracted total RNA samples at stages 53 and 60 were further digested with RNase-free DNase I (QIAGEN) and purified using a NucleoSpin RNA Clean-up kit (MACHEREY-NAGEL) for RNA-seq analysis.

### RT-PCR

Total RNA (100 ng for stage 53, 500 ng for stage 60, and 1 µg for stages 62 and 66) in 5.5 µL was reverse-transcribed by SuperScript III (Invitrogen) using random primer (15mer) in a 10 µL scale or in a proportionally scaled volume. PCR was performed using 1 µL of the obtained cDNA for 32 cycles. *gapdh* was used as an internal standard. Gene-specific primers were designed to span an intron, if any (Table S1).

### RNA-seq analysis

Isolated total RNAs were treated with DNase I and purified as mentioned above. Next-generation sequencing libraries were generated using the TruSeq Stranded mRNA Sample Preparation kit (Illumina). The libraries were subjected to single-end sequencing of 81 bp fragments using the Illumina NextSeq 550 platform. The *X. laevis* genome sequence and the gene models used for the read mapping were XENLA_9.2_genome.fa and XENLA_9.2_Xenbase_longest.gff3, which were downloaded from Xenbase (http://www.xenbase.org/entry/). Trim Galore! (version 0.6.6), HISAT2 (version 2.2.1), and StringTie (version 2.1.4) were used for the preprocessing, mapping, and read count, respectively (Kim et al., 2019; Pertea et al., 2016). The public RNA-seq datasets of the adult stomach and intestine (Session et al., 2016) were similarly processed. Normalization of read counts and identification of differentially expressed genes (DEGs) were performed using an *R* package, *DESeq2*. Genes with adjusted p-value (padj) < 0.05 and a minimum 2-fold expression ratio were defined as DEGs. Principal component analysis (PCA) plot, hierarchical-clustering heatmap, and the volcano plot were drawn using *R* packages *DESeq2*, *pheatmap*, and *EnhancedVolcano*, respectively. Gene ontology (GO) terms for each gene were annotated using eggNOG-mapper v2 (Cantalapiedra et al., 2021) (Table S3), and GO enrichment analysis for DEGs was performed using clusterProfiler (Wu et al., 2021). To produce a comprehensive list of transcription factors (TFs) from the *X. laevis* gene models, TBLASTN against *X. laevis* nucleotide sequences was performed using 1,237 *X. tropicalis* TFs listed by Blitz et al., (2017) as queries, and the top two hits were defined as putative orthologs. Reciprocal BLAST hits were also incorporated into the list, but genes that reside on chromosomes different from those of the corresponding *X. tropicalis* genes were excluded. In addition, MOTIF Search in GenomeNet (https://www.genome.jp/tools/motif/) was used to exclude genes which do not have any DNA binding motif. If necessary, gene synteny and sequence alignments were manually confirmed. Consequently, a total of 2,435 TFs (including L and S homeologs) were identified. They were hierarchically clustered according to their expression patterns using the cutree function in *R*. The number of the clusters was empirically determined to be 50. Z-scored transcripts per million (TPM) was used for the unit of gene expression level in the clustering analysis. Clustering results were visualized by *heatmap.2* package in *R*.

## Results

### 1. Histological analysis of pylorus and pyloric sphincter formation

To clarify when the stomach body, antrum, pylorus region, and duodenum are regionalized, and to determine when and where the pyloric sphincter is formed, we first histologically examined the stomach-duodenum region of embryos (stage 42/43), pre-metamorphic tadpoles (stages 51 and 53), metamorphic climax tadpoles (stages 60, 61, and 63), and froglet (stage 66) as well as adults.

#### 1-1 Embryonic and pre-metamorphic stages

To demarcate the stomach-duodenum region at embryo and tadpole stages, externally identifiable landmarks were defined and named as follows (Fig. S1): the anterior constriction, which is located between the stomach body and posterior stomach (magenta arrowheads in Fig. S1D,E); the flexure in the posterior stomach (magenta asterisks in Fig. S1A,B,D-F), which is described as the gastroduodenal (GD) loop by Bloom et al. (2013); and the posterior constriction (magenta arrows in Fig. S1D,E), which is followed posteriorly by the bile duct opening (white arrowheads in Fig. S1D,E), a characteristic of the duodenum.

At embryonic stage 42/43, the endodermal (en) and mesodermal (me) layers (distinguished by the amount of yolk granules) of the posterior stomach (p.st) and those of the duodenum (du) were morphologically uniform around the pancreas (pa) (Fig. 1A,A1). At tadpole stages (stages 46-53), the posterior stomach and duodenum dramatically elongated compared to the embryonic stages (Fig. S1B-D). The stomach body (st.b) and the posterior stomach (p.st) were distinguishable from their external appearance: the stomach body appeared brownish while the posterior stomach was lighter in color (Fig. S1D).

**Fig. 1.**
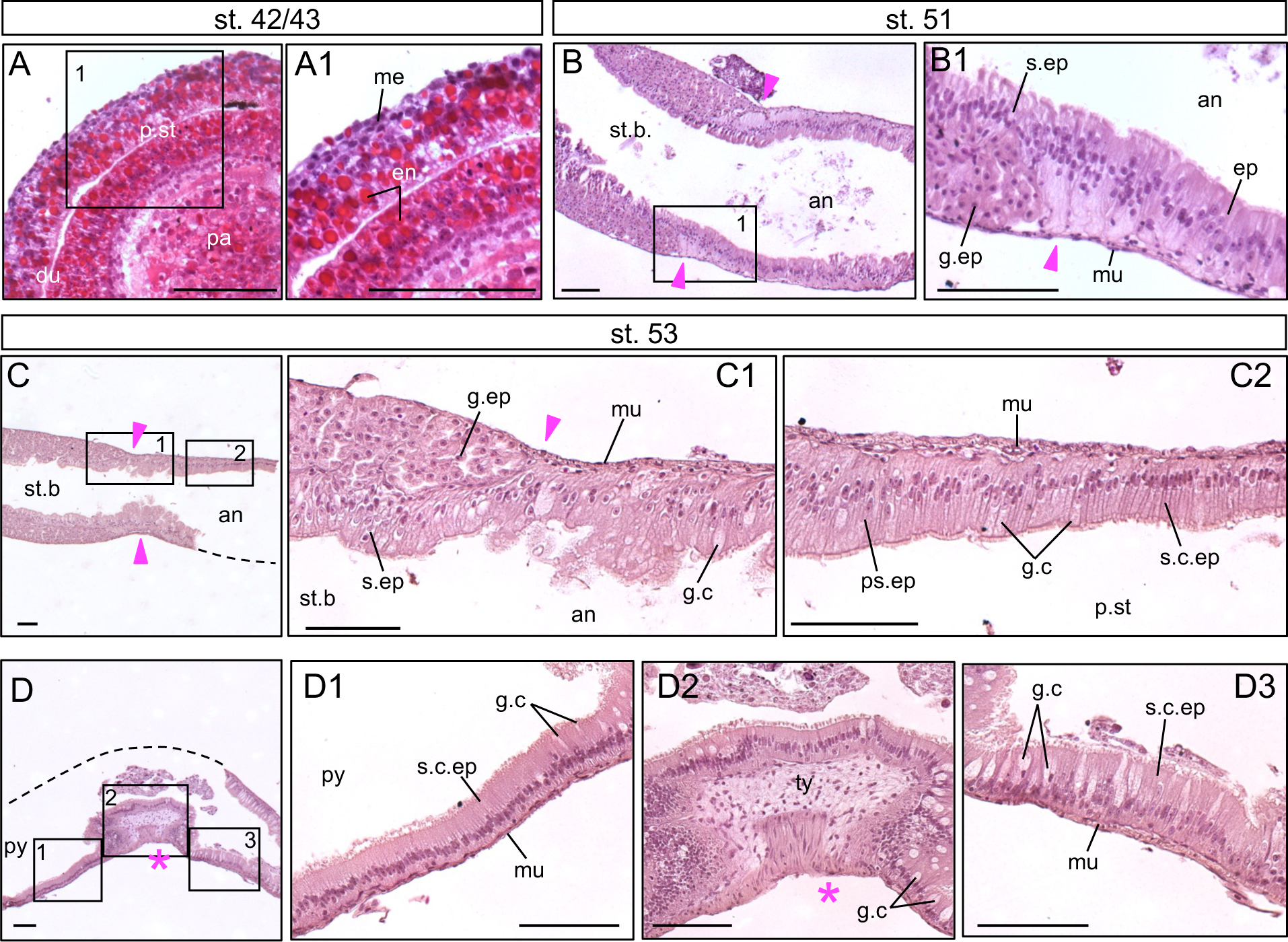
Histology of the stomach-duodenum region at embryonic to larval stages. Hematoxylin-eosin-stained sagittal sections of the stomach-duodenum region at stage 42/43 (A), stage 51 (B), stage 53 (C, D) in *Xenopus laevis*. Squares with nos. 1 - 3 in (A-D) are magnified in the corresponding panels A1-D3. As yolk granules were strongly stained in red with eosin (A), the endodermal layer is recognized as a yolk-rich region, compared to the mesodermal layer containing less yolk. Panel D is flipped to orient in the same direction as the others (anterior to the left). Dashed lines in (C, D) indicate the outlines of missing parts. Magenta asterisk, the pyloric flexure (see Discussion). Magenta arrowhead, anterior constriction. an, antrum; du, duodenum; en, endoderm; ep, epithelium; g.c, goblet cell; g.ep, glandular epithelium; me, mesoderm; mu, muscle; pa, pancreas; p.st, posterior stomach; ps.ep, pseudostratified epithelium; py, pylorus region; s.ep, surface epithelium; s.c.ep, simple columnar epithelium; st.b, stomach body; ty, typhlosole. Scale bars, 100 μm except for 50 μm in A1.

To search for the boundary between the pylorus region and duodenum at pre-metamorphic tadpole stages, we observed sections of the region between the stomach body and duodenum at stages 51 and 53 (Fig. S1D for external morphology). At stage 51, the epithelium of the stomach body consisted of the surface epithelium (s.ep) and the underlying glandular epithelium (g.ep) (Fig. 1B,B1), as reported (Lametschwandtner et al., 2012). From anterior to posterior, the epithelial nuclei tended to position more basally and the height of the cells became shorter, as has been reported at stage 48 (Smith et al., 2000a). Similar to stage 51, at stage 53, the surface and glandular epithelia (s.ep and g.ep) of the stomach body were observed rostral to the anterior constriction (Fig. 1C,1C1,D). In the antrum caudal to the anterior constriction, pseudostratified epithelium continued from the surface epithelium of the stomach body (Fig. 1C1), but at more caudal positions, the epithelium gradually turned into simple columnar epithelium with scattered goblet cells (g.c) (Fig. 1C1,C2). Further caudally, the epithelium became thinner with their nuclei positioned closer to the basal side (Fig. 1C2). The muscle layers (mu) of the stomach body and posterior stomach were uniformly thin (Fig. 1C1,C2). The simple columnar epithelium with goblet cells continued caudally (Fig. 1D). Reportedly, goblet cells are present in the adult intestinal epithelium and also in the transitional zone that is between the stomach body and the duodenum in the tadpole of *Rana* (Bodegas et al., 1997) and *Xenopus* (Chalmers and Slack, 1998; Chalmers et al., 2000). More caudally, the typhlosole (ty) and many goblet cells were observed (Fig. 1D1-3). The typhlosole is postulated to be the characteristic fold in the anterior small intestine of *X. laevis* at pre-metamorphic tadpole stages (Marshall and Dixon, 1978; Sterling et al., 2012). However, according to a previous report by Ishizuya-Oka et al. (Ishizuya-Oka et al., 1997), a typhlosole-like structure also appears to be present in the region rostral to the intestine. In addition, this region is negative in immunostaining for the intestinal marker IFABP (intestinal fatty acid binding protein) (Ishizuya-Oka et al., 1997). These data suggest that the typhlosole is present not only in the intestine but also in the pylorus region. This notion is further supported by the description by Nieuwkoop and Faber (1967) (Nieuwkoop and Faber, 1967) “typhlosole formed in duodenum and marginal gut at stage 51,” in which “marginal gut” most likely corresponds to the pylorus region. In conclusion, the pylorus region has some intestinal features and corresponds to the posterior portion of the transitional zone. Throughout embryonic and pre-metamorphic stages, there is no clear epithelial or mesenchymal boundary between the pylorus region (or transitional zone) and the duodenum, nor is there any indication of sphincter formation (Fig. 1D1).

**Fig. 2.**
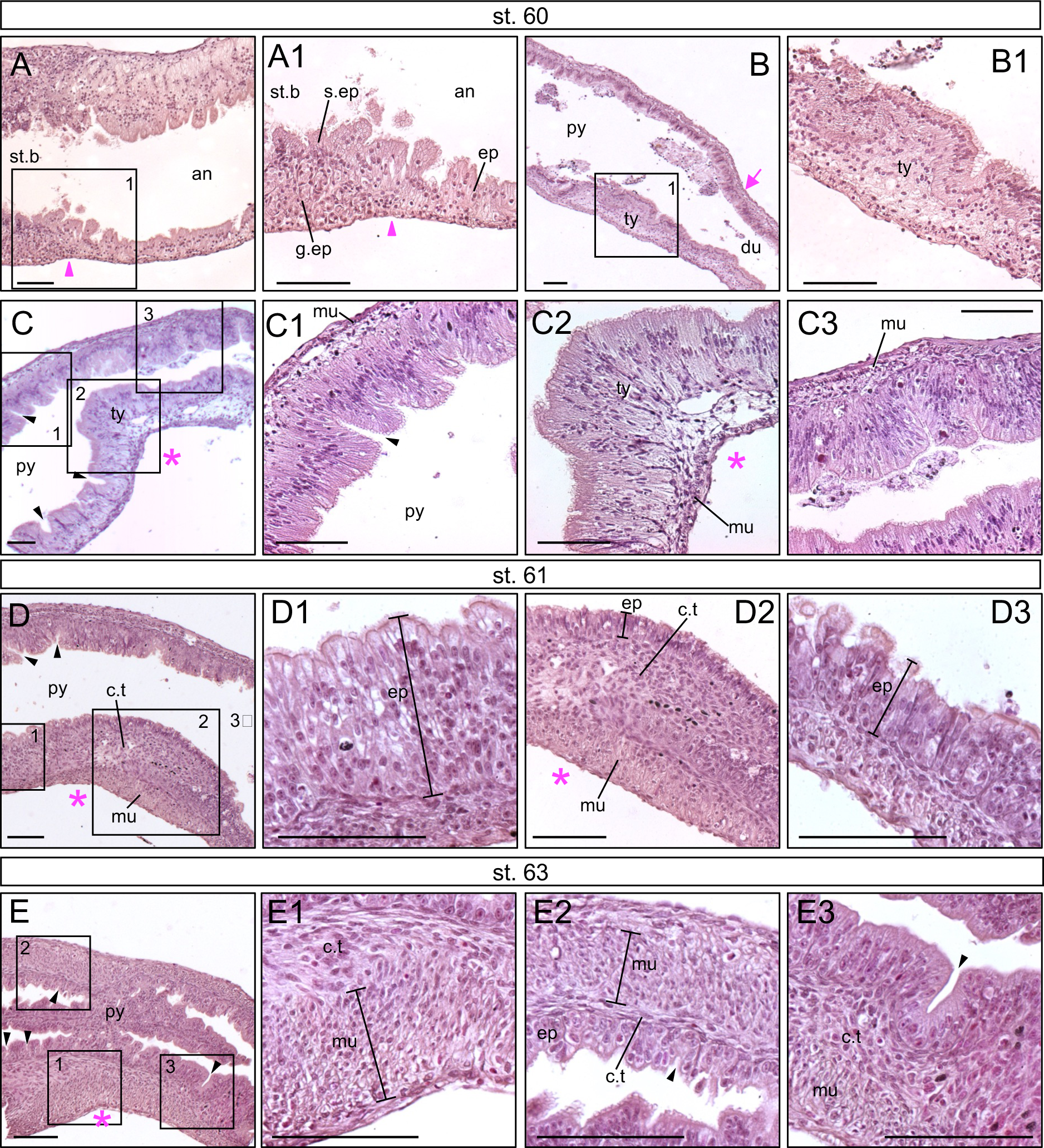
Histology of the stomach-duodenum boundary region during metamorphosis. Sagittal sections of the stomach-intestine boundary region at stages 60 (A, B, C), 61 (D), and 63 (E). Squares with nos. 1–3 in (A–E) are magnified in the corresponding panels A1–E3. Panels D and E are flipped to orient in the same direction as A and B (anterior to the left). Panels C1, D1 and D3 are located partially or completely outside of panels C and D (see Fig. S2 for the position of squares 1 and 3 in D). Magenta asterisk, the pyloric flexure (see Discussion). Magenta arrow in (B), posterior constriction. Black arrowhead, pyloric gland; magenta arrowhead, anterior constriction. an, antrum; c.t, connective tissue; du, duodenum; ep, epithelium; g.ep, glandular epithelium; mu, muscle; py, pylorus region; s.ep, surface epithelium; st.b, stomach body; ty, typhlosole. Scale bars, 100 μm.

**Fig. 3.**
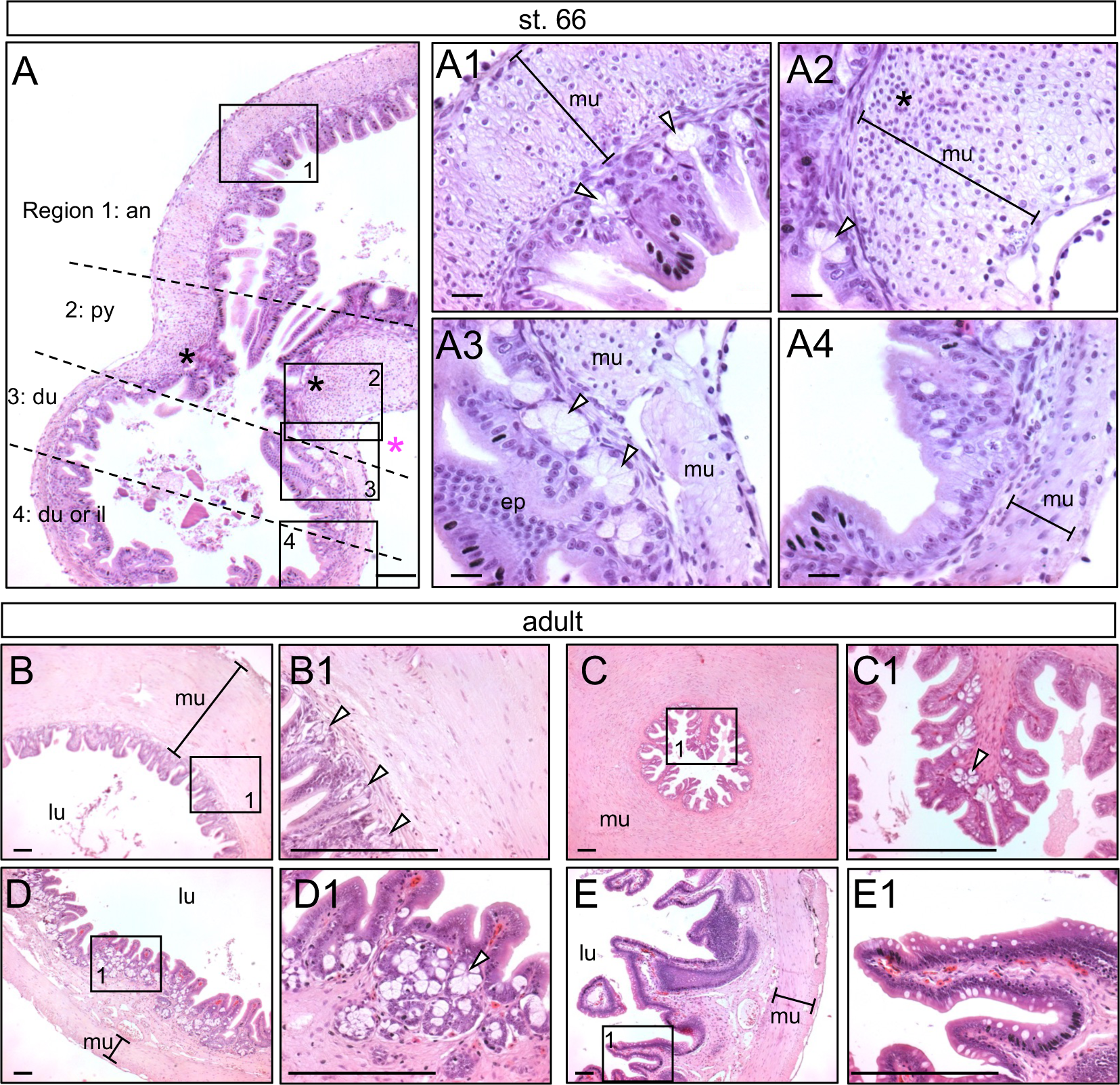
Histology of the stomach-duodenum boundary region after metamorphosis. (A) Sagittal sections at stage 66. The stomach-duodenum boundary region is divided into Regions 1-4 with dashed lines (see the text). Squares with nos. 1-4 are magnified in A1-A4. (B-E) Cross sections of the adult GI tract at four positions. Panels B-E correspond to Regions 1 to 4 in (A). Square 1 in (B-E) is magnified in B1-E1. Black asterisks in (A, A2) indicate the inner and outer nuclear-dense regions of the muscle thickening in the pyloric sphincter at the pyloric flexure (magenta asterisk), which overlap with the posterior constriction at this stage. White arrowhead, mucus gland. an, antrum; du, duodenum; ep, epithelium; il, ileum; lu, lumen; mu, muscle; py, pylorus region. Scale bars, 20 μm in (A1-A4) and 100 μm in the others.

#### 1-2 Metamorphic stages

To determine the timing and location of pyloric sphincter formation, we examined sagittal sections of the stomach body-to-duodenum region at metamorphic-climax stages 60, 61 and 63. At stage 60, anterior and posterior constrictions of the GI tract were observed corresponding to those of stage 53 (Fig. S1E). At the anterior constriction, the surface and glandular epithelia of the stomach body were followed caudally by the pseudostratified epithelium of the antrum (Fig. 2A). The typhlosole (ty) was observed rostral to the posterior constriction (Fig. 2B,2B1). The epithelium rostral to the flexure (magenta asterisk) had many pits consisting of tall columnar cells (Fig. 2C; black arrowhead), probably forming pyloric glands, a marker of the posterior stomach. A bundle of circular muscle was still not observed at stage 60 in the pylorus region (Fig. 2C,C2).

At stage 61, the muscle layer (mu) in the inner curvature of the flexure became thickened in the connective tissue (c.t) of the typhlosole-like structure (Figs. 2D,D2, S2A,B), probably indicating the onset of sphincter formation. The inner curvature of the flexure corresponds to the dorsal side of the GI tract at the embryonic stage, because the pancreatic rudiment is attached there (Fig. S1) (Womble et al., 2016). It was reported that there is abrupt transition in epithelial morphology from the sphincter to duodenum in adult *Xenopus* (Smith et al., 2000), but such epithelial transition near the muscle thickening was not recognized (Figs. 2D3, S2). These observations suggest that sphincter formation begins on the inner (dorsal) side of the flexure within the pylorus region at stage 61 or earlier, before epithelial transition between the pylorus region and duodenum is formed.

At stage 63, the stomach anterior to the flexure was enlarged (Fig. S1F). The inner circular muscle layer at the flexure was further thickened with high nuclear density (Fig. 2E,E1), compared to stage 61 (Fig. 2D,D2). Thickening of the muscle was also observed on the outer (ventral) side of the flexure (Fig. 2E,E2), indicating that the pyloric sphincter is being formed. The epithelium around the muscle thickening had pyloric pits (Fig. 2E,E3; black arrowheads), suggesting that the sphincter is formed within the pylorus region rather than at the posterior end of the stomach.

#### 1-3 Froglets and adults

We next examined the positional relationship between the epithelial boundary and the pyloric sphincter at the froglet (stage 66) and adult stages. At stage 66, a constriction at the stomach-duodenum boundary was externally identified (Fig. 3A). Based on the epithelial structure and the thickness of the muscle layer, we determined the following four regions around the constriction (Fig. 3A). Region 1 (the antrum) had epithelium with mucus glands and a thick circular muscle layer (Fig. 3A1). Region 2 (the pylorus region) had mucus glands (white arrowheads) and the sphincter (black asterisks) on both inner and outer curve sides of the flexure (Fig. 3A). The sphincter at the inner curve of the flexure was much thicker than on the outer side (Fig. 3A,3A2; black asterisks). The pylorus region in the froglet shrunk dramatically compared to that of stage-61 tadpoles (Figs. 2D, S2), and the flexure observed before stage 61 appeared to converge with the posterior constriction by stage 66. Region 3, the anterior part of the duodenum, had dense mucus glands in its epithelium, and its muscle layer (mu) was thinner than in Regions 1 and 2 (Fig. 3A3). The mucus glands in Regions 2 and 3 may correspond to the pyloric and duodenal glands in amniotes, respectively. These data indicate that clear boundaries in the epithelial and muscle layers of the stomach and duodenum are formed at the posterior end of the pylorus by the end of metamorphosis. Region 4 did not have the mucus glands but had a thin circular muscle layer (Fig. 3A4), and is likely to be the posterior duodenum or the ileum. These four regions were also observed in the adult frog, whose muscle layer of each region was much thicker than at stage 66 (Fig. 3B-E).

### 2. Expression analyses of pylorus/sphincter-related genes in the posterior stomach and duodenum

To examine the role of pylorus/sphincter-related genes in *Xenopus*, we analyzed developmental expression of *barx1*, *cdx2*, *nkx2-5*, *gata3*, *sox9*, *bmp4*, *nkx3-2* and *grem1*. We performed whole-mount in situ hybridization (WISH) for embryonic (stage 42/43) and early tadpole stages (stage 46/47), whereas section in situ hybridization (ISH) and RT-PCR were used for the late tadpole (stage 53), metamorphic (stages 60-62), and froglet (stage 66) stages due to strong non-specific signals in WISH after stage 48.

#### 2-1 barx1 and cdx2

*barx1* and *cdx2* are the regional markers for the stomach and intestine, respectively, in mice (Gao et al., 2009; Kim et al., 2005). In the tailbud embryo (stage 42/43), *barx1* was specifically expressed in the stomach with a clear posterior boundary (Fig. 4A). At the early tadpole stage (stage 46), the posterior boundary of *barx1* expression coincided with the posterior constriction (Fig. 4C,C1), suggesting that the posterior constriction is the posterior end of the stomach. In contrast to a previous study (Chalmers et al., 2000), *cdx2* expression was detected in the flexure and posterior intestine at stage42/43, and was more broadly expressed from the stomach to the intestine at stage 46 (Fig. 4B,D). The anterior expression boundary of *cdx2* was not clearly defined, and apparently overlapped with the *barx1* expression domain in the stomach (Fig. 4A,C), indicating that there is no clear expression boundary between *barx1* and *cdx2* at the embryonic and tadpole stages. This data suggests that the molecular boundary of the stomach and intestine in *X. laevis* is not yet established before metamorphosis.

**Fig. 4.**
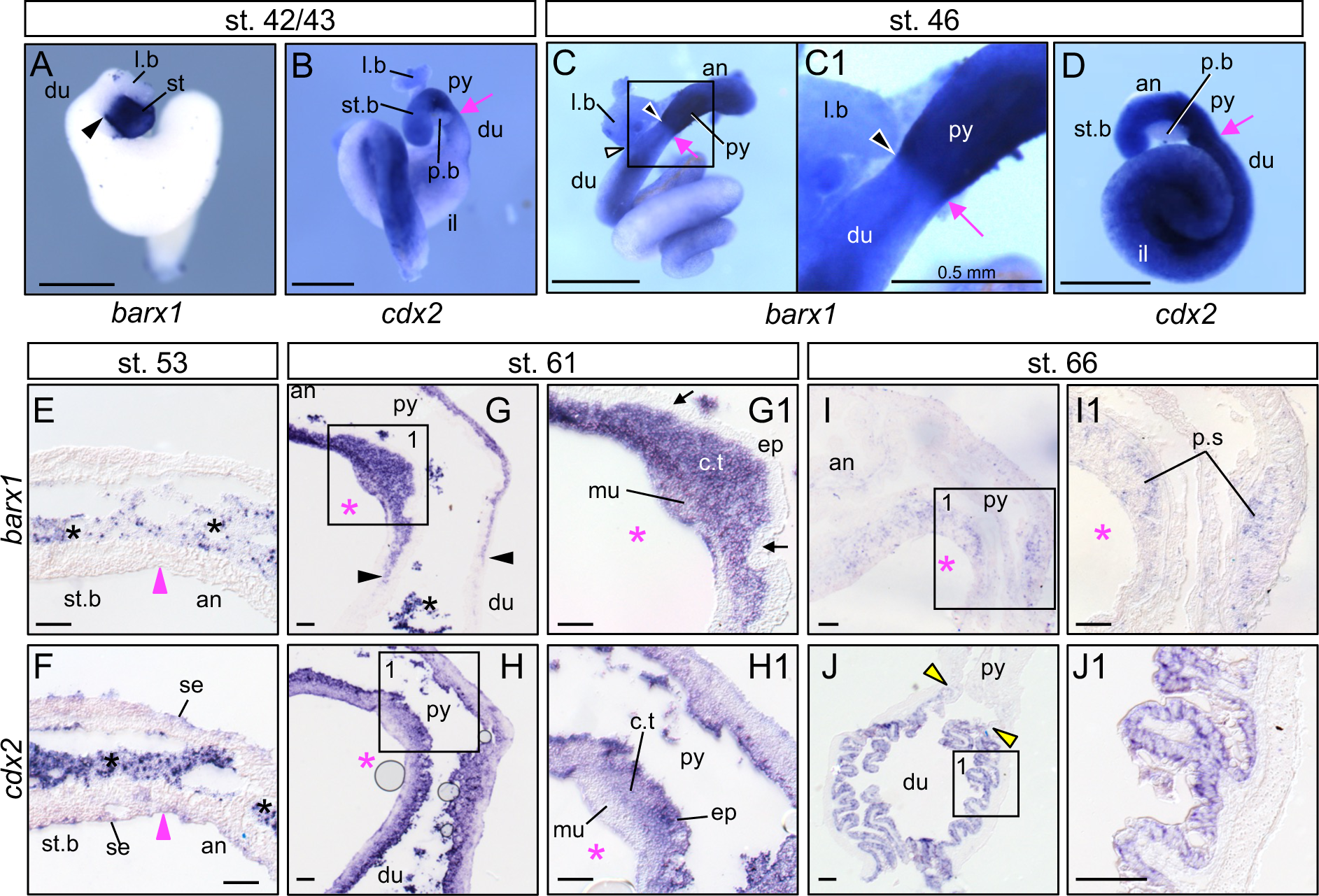
Expression patterns of *barx1* and *cdx2* in the stomach-duodenum region at late tailbud to froglet stages. (A-D) Whole-mount in situ hybridization of isolated GI tracts for *barx1.L* (A,C) and *cdx2.L* (B, D) at stages 42/43 (A, B) and 46 (C, D). *barx1* staining in the stomach at stage 46 was strongly detected with the posterior boundary, but relatively weak staining in the duodenum might be nonspecific (C). (E-J) Section in situ hybridization of the GI tract for *barx1.L* and *cdx2.L* around the anterior constriction at stage 53 (E, F) and around the pyloric flexure at stage 61 (G, H) and stage 66 (I, J). Squares with no. 1 in (G-J) are magnified in panels G1-J1. Black asterisks in (E,F) indicate nonspecific signal for undigested contents; bubbles in (H) are artifacts; weak signal of serosa (se) with *cdx2* probe in (F) is probably nonspecific staining. Magenta asterisk, pyloric flexure. Black arrow (G1), lack of in situ signal; magenta arrow, posterior constriction;. Black arrowhead (G), posterior boundary of *barx1* expression; magenta arrowhead (E,F), anterior constriction between the stomach body (st.b) and antrum (an); white arrowhead (C), the bile duct opening; yellow arrowhead (J), the anterior boundary of *cdx2* expression. an, antrum; c.t, connective tissue; du, duodenum; ep, epithelium; il, ileum; l.b, liver bud; mu, muscle; p.b, pancreatic bud; p.s, pyloric sphincter; py, pylorus region; st, stomach; st.b, stomach body. Scale bars, 1 mm otherwise indicated in (A-D) and 100 μm in (E-J1).

At the late tadpole stage (stage 53), *barx1* expression was not detected by ISH in the stomach body and antrum even after chromogenic reaction for two days (Fig. 4E). This is not due to technical errors, because control staining of *acta2* mRNA with the same series of sections gave strong signals in the smooth muscle (Fig. S3) and the same *barx1* probe preparation worked with the stomach section of metamorphic tadpoles (see Fig. 4G). Similarly, *cdx2* expression was not detected in the stomach body and antrum (Fig. 4F), suggesting that *barx1* and *cdx2* are downregulated once the formation of the larval stomach is complete.

At the metamorphic climax (stage 61), *barx1* was strongly expressed again in the connective tissue and muscle layers of the antrum and pylorus region (Fig. 4G), but not in their epithelium (Fig. 4G1; black arrows; see Fig. 2D for morphology). Its expression had a clear posterior boundary caudal to the developing pyloric sphincter (p.s) (Fig. 4G; black arrowheads). *cdx2* was expressed in the epithelium of the antrum-to-duodenum region, weakly in the connective tissue, but barely in the muscle layer (Fig. 4H,H1).

At stage 66 (froglet), *barx1* expression was detected in the pyloric sphincter, but unlike during metamorphosis, it was hardly detected posterior to the sphincter (Fig. 4I,I1). In Region 3 (anterior duodenum), *barx1* expression was not detected (Fig. 4I), but the anterior expression boundary of *cdx2* was recognized (Fig. 4J,J1; yellow arrowheads). *cdx2* expression was mostly restricted to the epithelium of Regions 3 and 4 (duodenum; Fig. 4J). In summary, the expression domains of *barx1* and *cdx2* overlap in the stomach during embryonic to metamorphic climax stages but are clearly demarcated after metamorphosis. Since *barx1* is expressed more posteriorly to the pyloric sphincter at stage 61 (Fig. 4G), the sphincter is suggested to be formed within the posterior stomach, not at the boundary of the stomach and duodenum, consistent with our histological observation.

#### 2-2 sox9, bmp4, nkx3-2, gata3, grem1, and nkx2-5

We further analyzed expression of *sox9, bmp4, nkx3-2, gata3, grem1*, and *nkx2-5*, which are suggested to be involved in the formation of the pylorus and pyloric sphincter in amniotes (Li et al., 2009; Moniot et al., 2004; Self et al., 2009; Smith et al., 2000a; Smith et al., 2000b; Theodosiou and Tabin, 2005; Verzi et al., 2009). The expression domains of these pylorus/sphincter-related genes at stage 42/43 (late tailbud stage) were overall similar to amniotes as follows. *sox9* was expressed strongly in the stomach body (st.b) and pylorus region (py), but not in the antrum (an) (Fig. 5A), as has been implicated in mouse and chicken embryos (Theodosiou and Tabin, 2005; Udager et al., 2010). *bmp4* and *nkx3-2* were expressed in the pylorus region but weakly in the stomach body (Fig. 5B,C; (Smith et al., 2000b; Verzi et al., 2009). *gata3* was broadly expressed in the stomach (Fig. 5D; Li et al., 2009). *grem1* was broadly expressed around the pylorus region (Fig. 5E), whereas *nkx2-5* expression was more restricted to the pylorus region (Fig. 5F) as previously reported (Li et al., 2009; Moniot et al., 2004; Self et al., 2009; Smith et al., 2000a). Thus, despite the absence of the pyloric sphincter, various pylorus/sphincter-related genes are already expressed in the pylorus region of *Xenopus* embryos.

**Fig. 5.**
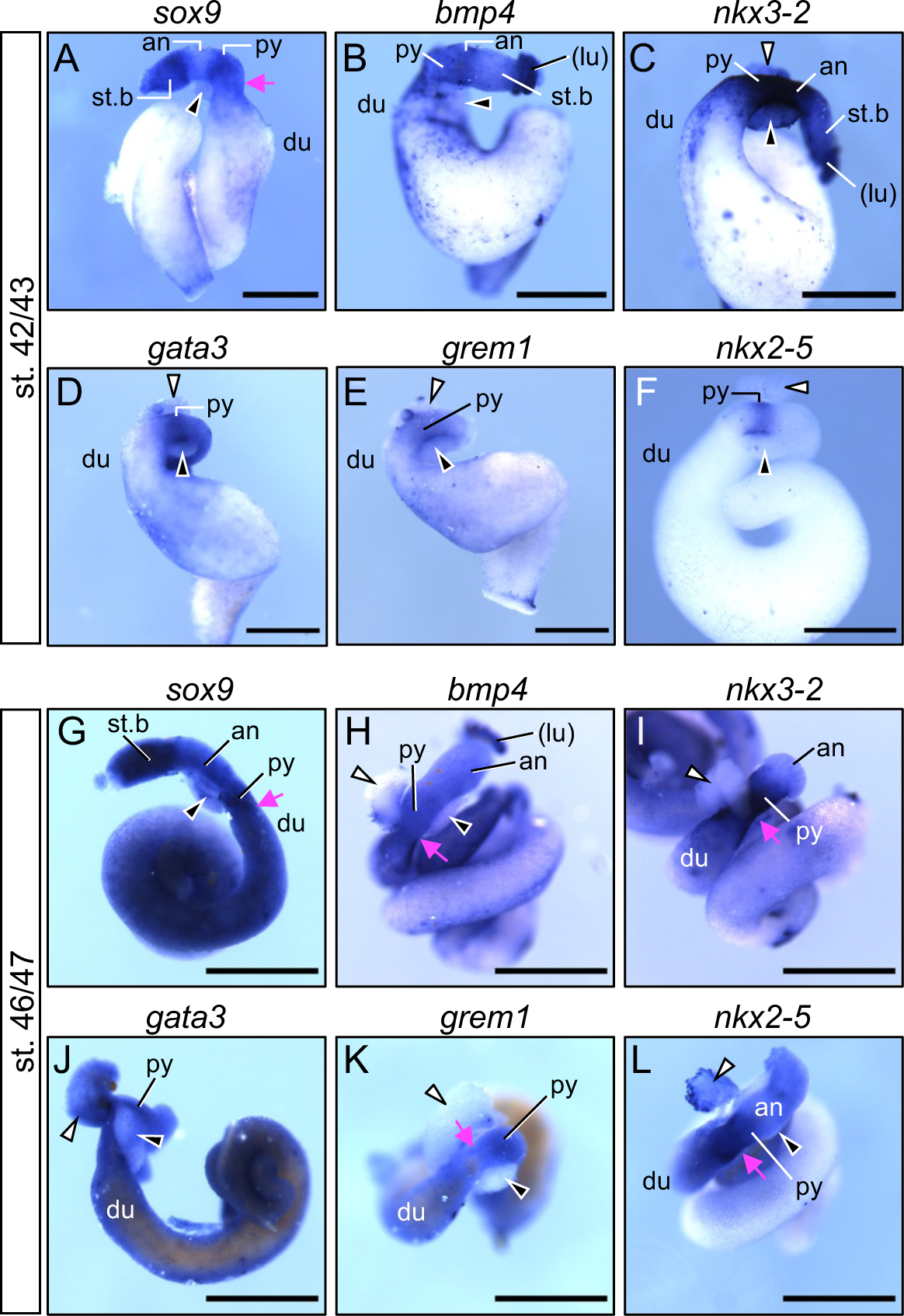
Whole-mount in situ hybridization of the GI tract for pylorus/sphincter-related genes. Whole-mount in situ hybridization of isolated GI tracts was performed for the pylorus/sphincter-related genes, *sox9, bmp4, nkx3-2, gata3, grem1*, and *nkx2-5*. (A-F) Late tailbud embryos (stage 42/43). (G-L) Early tadpoles (stage 46/47). Magenta arrow, posterior constriction. Black arrowhead, pancreatic bud; white arrowhead, liver bud. an, antrum; du, duodenum; (lu), unremoved part of lung; py, pylorus region; st.b, stomach body. Scale bars, 1 mm.

At the early-tadpole stage (stage 46/47), the pylorus/sphincter-related genes were in the stomach in a similar manner as stage 42/43, but their expression appeared to extend beyond the posterior constriction into the duodenum (Fig. 5G-L). The antrum marked by weak *sox9* expression appeared to be elongated (compare Fig. 5A and G), consistent with external appearance (Fig. S1). By contrast, at the late-tadpole stage (stage 53), *sox9*, *gata3*, and *nkx2-5* were barely expressed in the stomach body and antrum (Fig. 6A-C), suggesting that their expression is restricted to the developing GI tract like *barx1* (Fig. 4E).

**Fig. 6.**
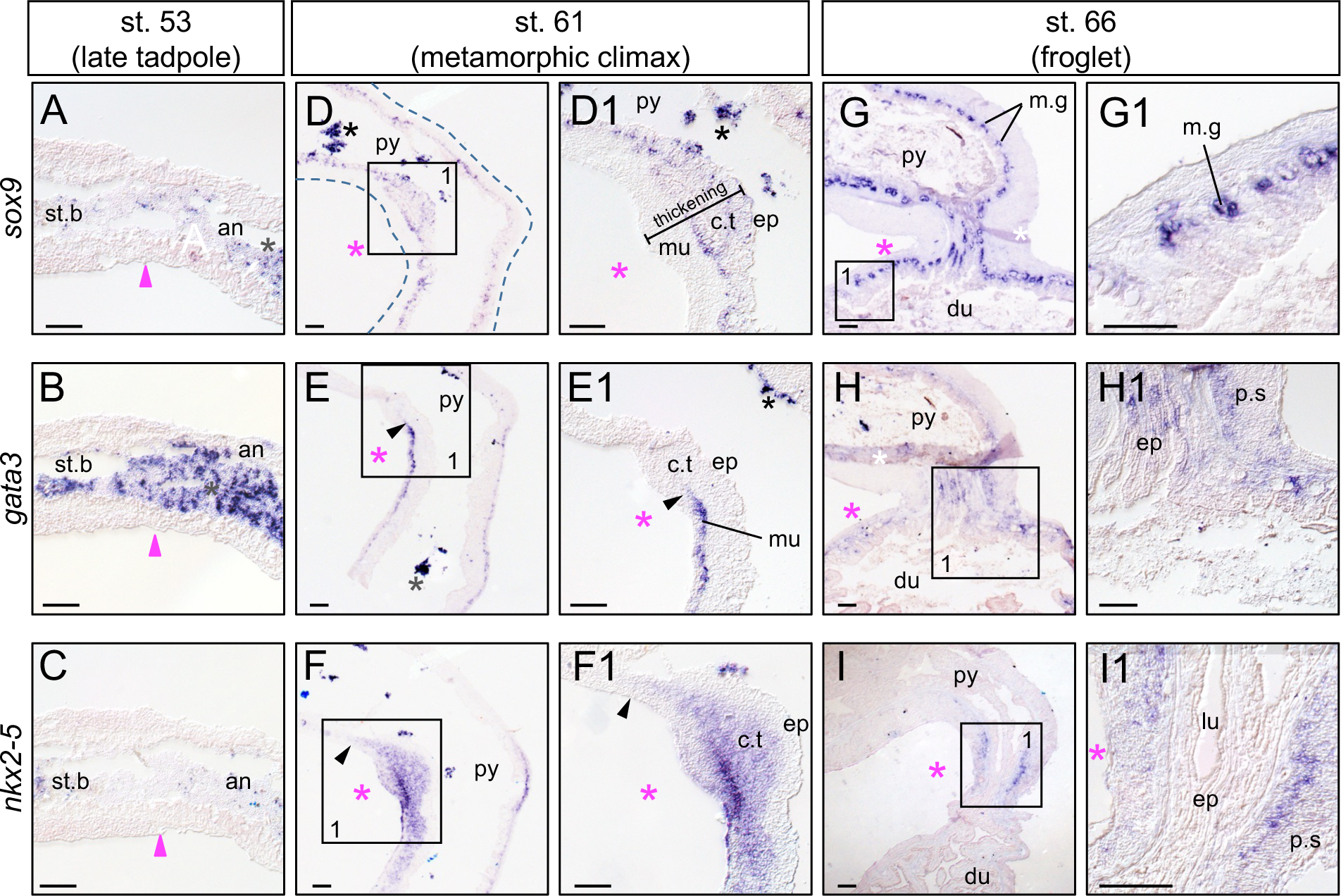
Section in situ hybridization of the stomach-duodenum region for pylorus/sphincter-related genes. Section in situ hybridization of the stomach-duodenum region was performed for the pylorus/sphincter-related genes, *sox9, gata3* and *nkx2-5*. (A-C) The stomach region around the anterior constriction at stage 53. (D-I) The stomach-duodenum boundary region at stage 61 (D-F) and stage 66 (G-I). Squares in (D–I) are enlarged in panels D1-D1-I1. Dashed lines in (D) indicates the contour of the section. Black asterisks indicate nonspecific signal for undigested contents. White asterisks in (G, H) indicates artifact of section. Magenta asterisk, the pyloric flexure, Black arrowhead, anterior boundary of expression; magenta arrowhead, the anterior constriction. an, antrum; c.t, connective tissue du, duodenum; ep, epithelium; lu, lumen; m.g, mucus gland; mu, muscle; p.s, pyloric sphincter; py, pylorus region; st.b, stomach body. Scale bars, 100 μm.

At metamorphic climax (stage 61), *sox9*, *gata3*, and *nkx2-5* were expressed again (Fig. 6D-F), implying their functions in the stomach remodeling. *sox9* was expressed in the epithelium (ep) and connective tissue (c.t) but not in the muscle layer (mu) of the flexure (Fig. 6D). *gata3* was expressed in the muscle layer of the flexure-to-duodenum region and had an anterior expression boundary (Fig. 6E,E1; black arrowhead). *nkx2-5* was expressed in the connective tissue and muscle layers in and posterior to the flexure (Fig. 6F), overlapping with *barx1* (Fig. 4G). Thus, it is likely that *sox9*, *gata3*, and *nkx2-5* are involved in the pyloric sphincter formation, as in amniotes (Li et al., 2014; Prakash et al., 2014; Theodosiou and Tabin, 2005; Verzi et al., 2009).

At the froglet stage (stage 66), *sox9* expression was not detected in the pyloric sphincter (p.s), but in the mucus glands (m.g) of the pylorus and duodenum (du) (Fig. 6G,G1), suggesting that *sox9* has a new role in differentiated cells after metamorphosis. *gata3* expression was weakly detected in the epithelial layers as well as the luminal side of pyloric sphincter (Fig. 6H,H1), where *nkx2-5* was also weakly expressed (Fig. 6I,I1), implying their functions other than sphincter formation after metamorphosis.

#### 2-3 RT-PCR analysis of the antrum and pylorus regions before and during metamorphosis

To validate the results of ISH, we performed semiquantitative RT-PCR analysis. GI tracts at pre-metamorphic and metamorphic tadpole stages (stages 53 and 60, respectively) were dissected into 1-mm pieces using the anterior constriction as a positional marker (Fig. 7A). The pieces were numbered from anterior to posterior with prefixes T (tadpole stage) or M (metamorphic stage). For metamorphic and froglet stages (stages 62 and 66, respectively), the flexure was used as the positional marker to dissect the GI tract into regions 1 (antrum), 2 (around the pylorus region), and 3 (duodenum) (Fig. 7A).

**Fig. 7.**
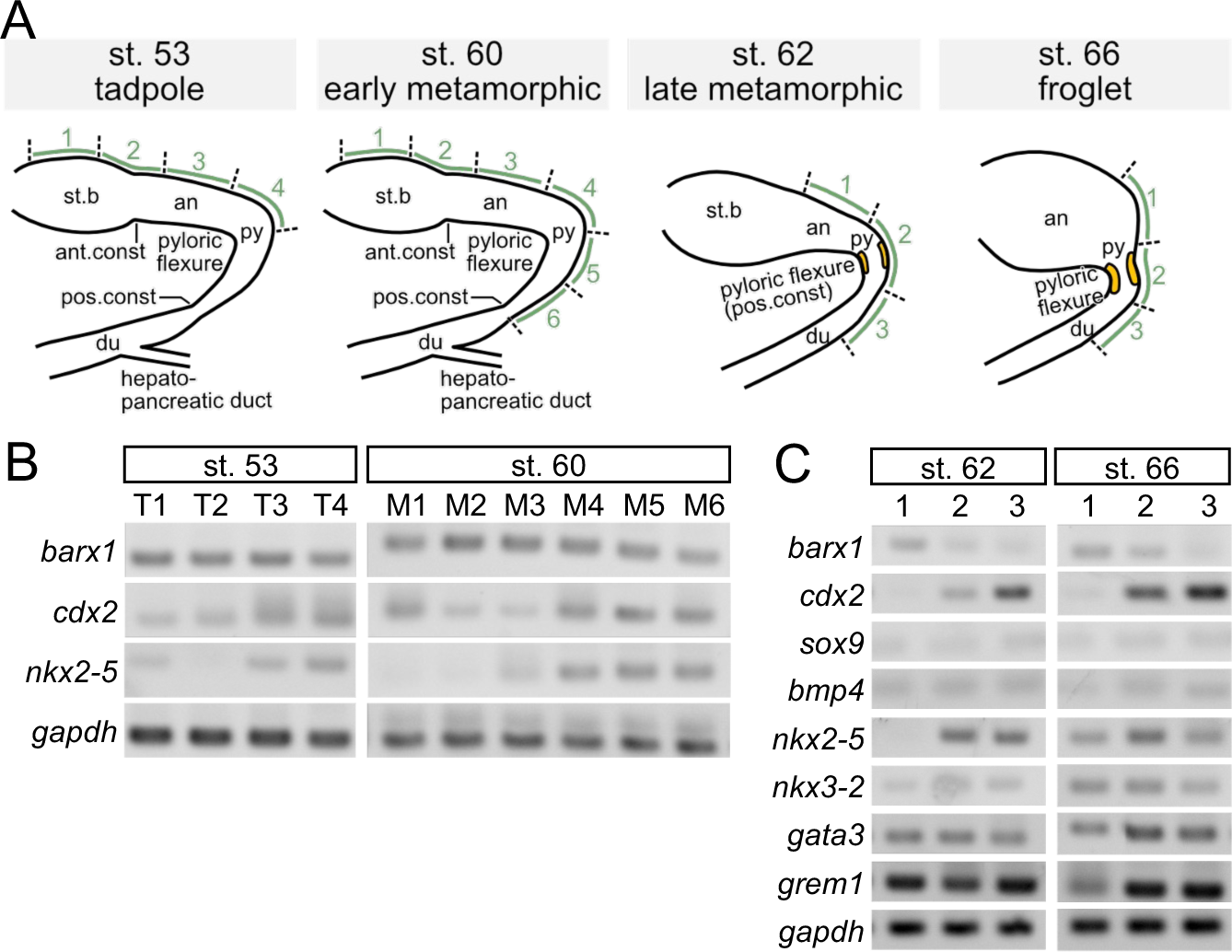
RT-PCR analyses of pylorus/sphincter-related genes in the stomach-duodenum region. (A) Positions of dissected GI tracts for total RNA extraction at stages 53, 60, 62, and 66. GI tracts were dissected into T1 to T4 for pre-metamorphic tadpoles (stage 53) and M1 to M6 for early metamorphic climax tadpoles (stage 60), regions 1 to 3 for late metamorphic tadpole (stage 62) and froglets (stage 66). (B, C) Semiquantitative RT-PCR at stages 53 and 60 (B) and stages 62 and 66 (C). *gapdh*, internal control for total RNA. T1 and M1, the posterior part of the stomach body; T2 and M2, the region across the anterior constriction; T3 and M3, the antrum; T4, M4 and M5, the pyloric flexure; M6, the region anterior to the posterior constriction. an, antrum; ant.const, anterior constriction; du, duodenum; pos. const, posterior constriction; py, pylorus region; st.b, stomach body.

At stage 53, *barx1* was expressed in T1 to T4 at a similar level (Fig. 7B, left panel). As *barx1* expression was barely detected by ISH (Fig. 4E), *barx1* could be broadly and weakly expressed in the stomach at pre-metamorphic stages. *cdx2* was expressed in T3 and T4 and weakly in T1 and T2 (Fig. 7B, left panel), suggesting a posterior-to-anterior expression gradient in the stomach, in contrast to the previous reports (Chalmers et al., 2000; Reece-Hoyes et al., 2002). *nkx2-5* was expressed in T1, T3, and T4, but barely in T2 (Fig. 7B, left panel). At the metamorphic climax (stage 60), *barx1* and *cdx2* were both expressed in M1 to M6, whereas *nkx2-5* was more specifically expressed in M3 to M6 (Fig. 7B, right panel). The absence of *nkx2-5* expression in M1 and M2 indicates that the expression of *cdx2* in M1 and M2 is not due to contamination of the posterior portion, further supporting *cdx2* expression in the stomach before metamorphosis.

At the late metamorphic tadpole and froglet stages (stages 62 and 66), region 1 (antrum) expressed only *barx1*, region 2 (pylorus) expressed both *barx1* and *cdx2*, and region 3 (duodenum) expressed *cdx2*, but very weakly *barx1* (Fig. 7C), in consistent with the section ISH at stage 66 (Fig. 4I, J). By contrast, expression of pylorus/sphincter-related genes, *sox9*, *bmp4*, *nkx2-5*, *nkx3-2*, *gata3* and *grem1*, was broadly detected from the posterior stomach to the duodenum at stage 66 (Fig. 7C), implying that their roles are changed from sphincter formation to others after metamorphosis, as exemplified by *sox9* expression in the mucus glands (Fig. 6G).

### 3. RNA-seq analysis of the stomach body, antrum, and pylorus regions at pre-metamorphic and metamorphic climax stages

To search for uncharacterized genes that could be involved in the pylorus and sphincter formation, especially those encoding transcription factors (TFs) and cell-cell signaling factors, we performed RNA-seq analysis on the stomach at pre-metamorphic (stage 53) and metamorphic climax (stage 60) stages. Four dissected stomach regions (T1 to T4 at stage 53 and M1 to M4 at stage 60; see Figs. 7A, S4) were designated as A, M, P, and PY for anterior, mid, posterior, and pylorus regions at each stage (Fig. 8A). We prepared samples from three clutches of tadpoles at each stage, indicated by the suffixes _1, _2, and _3. For example, st53A_1 stands for the anterior region of clutch 1 tadpoles at stage 53. st53/st60A, st53/st60M, and st53/st60P samples were collected from three clutches; the st53PY sample was from clutch 3 (st53PY_3): and st60PY samples were from clutches 2 and 3 (st60PY_2 and _3).

**Fig. 8.**
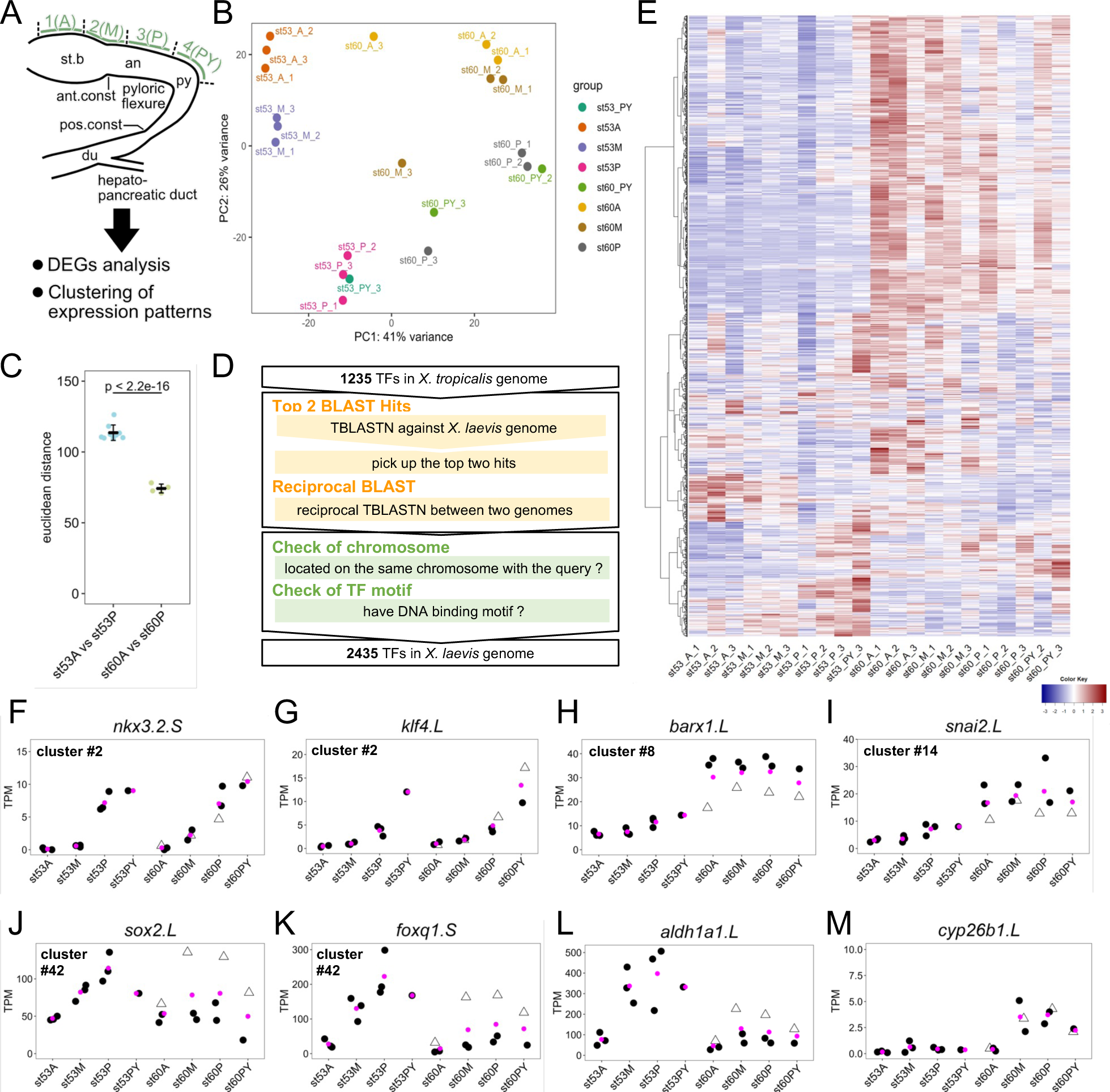
Regional RNA-seq analysis of stomach isolated from pre-metamorphic and metamorphic climax tadpoles. (A) Scheme of RNA-seq experiments. Stomach samples from pre-metamorphic (stage 53) and metamorphic climax (stage 60) tadpoles were dissected into A (anterior), M (mid), P (posterior), and PY (pyloric) regions. For example, st53A_1 indicates stage 53, anterior part, replicate #1. A, M, P, and PY correspond to the stomach body, the region containing the anterior constriction, the antrum, and the pylorus region, respectively. RNA-seq data were subjected to differential gene expression analysis and clustering of expression patterns. an, antrum; ant.const, anterior constriction; du; duodenum; pos.const, posterior constriction; py, pylorus region; st.b, stomach body. (B) Principal component analysis (PCA) plot of all samples. PC1 and PC2, principal components 1 and 2. (C) Comparison of anterior-posterior transcriptomic distances at stages 53 and 60. Euclidean distances between the st53A and st53P transcriptomes and between the st60A and st60P transcriptomes were calculated and plotted (see Fig. S5A). Brunner-Munzel test was used to calculate p-value. (D) Scheme of building a list of *X. laevis* transcription factors (TFs) using the list of *X. tropicalis* TFs (Blitz et al., 2017). 2,435 TFs including L and S homeologs were identified (Table S4), and 1881 TFs were annotated with gene names, whereas the other TFs only had either Xelaev IDs (74 TFs) or LOC IDs (480 TFs). Out of the 1,235 *X. tropicalis* TFs, 990 TFs (80%) had two or more homeologs in *X. laevis*, whereas 161 TFs (13%) only had either L or S homeolog. No orthologs were found for the remaining 84 TFs (7%). (E) Hierarchical clustering of expression profiles of TF genes. (F-M) Dot plots for expression levels of representative TFs in Groups 1 (F, G), 2 (H, I) and 3 (J,K), and genes involved in retinoic-acid pathways (L,M). (F) *nkx3-2.S*, (G) *klf4.L*, (H) *barx1.L*, (I) *snai2.L*, (J) *sox2.L*, (K) *foxq1.S*, (L) *aldh1a1.L*, and (M) *syp26b1.L*. Each plot is labeled with the group and cluster numbers for TF genes. The ordinate, transcripts per million (TPM). The sample names, st53A, st53M, st53P, st53PY, st60A, st60M, st60P, st60PY are indicated below the plot. Black dots indicate TPM values except for st60_3 (early) samples (triangles). Mean TPM values are shown with magenta dots.

Principal component analysis (PCA; Fig. 8B) and hierarchical clustering analysis (HCA; Fig. S5A) separated the dataset into replicate clusters, but all 4 samples, A, M, P, and PY, of st60 clutch 3 (referred to as st60_3 samples) did not cluster with the other two clutches of the same stage. In the PCA plot (Fig. 8B), the PC1 and PC2 axes are likely to represent the developmental stages and the anteroposterior positions, respectively. st60_3 samples were plotted between the st53 samples and the other st60 samples along the PC1 axis (Fig. 8B), suggesting that their actual stage could be earlier than st60_1 and _2. For convenience, st60_3 is referred to as st60_3 (early) hereafter. The Euclidean distance between st60A and st60P transcriptomes (74.21 ± 3.15) was significantly shorter than that between st53A and st53P transcriptomes (113.59 ± 5.45) (p-value < 2.2e-16 by the Brunner-Munzel test) (Figs. 8C, S5A). This indicates that the antero-posterior difference in gene-expression profiles is reduced during metamorphosis, possibly owing to up- or downregulation of thyroid-hormone (TH) responsive genes in the entire stomach and/or downregulation of position-specific genes (see below). Consistently, DEG (differentially expressed gene) analysis revealed that the numbers of DEGs between each two regions at stage 60 (Fig. S6D-F) were much smaller than at stage 53 (Fig. S6A-C). Regarding functional features of genes, gene ontology (GO) analysis on DEGs between st60P and st53P showed that GO terms related to developmental processes (“tube lumen cavitation,” “extracellular matrix disassembly,” etc.) and cell cycle progression (“DNA-dependent DNA replication,” “DNA packaging,” etc.) were enriched in st60P-upregulated gene sets (Fig. S5B; Table S3), whereas GO terms related to metabolic processes were enriched in the downregulated ones (Fig. S5C). st60P-upregulated DEGs included TH-inducible genes, such as *thibz.L/S, sox4.L/S*, and *shh.L* (Fig. S6G-I, underlined; see Figs. S8K,L and S9A), and st60P-downregulated DEGs included stomach-body-specific genes, such as *gastric H(+)-K(+)-ATPase subunit* genes, *atp4b.L/S* (Fig. S6H, magenta box; S6J,K). These tendencies are in good agreement with general features of metamorphosis, such as upregulation of genes associated with cell death, proliferation and differentiation, and downregulation of intracellular metabolic processes (Buchholz et al., 2007; Heimeier et al., 2010), supporting the validity of our datasets.

To focus on TF genes, we next built a comprehensive list of 2,435 TF genes (including L and S homeologs) in the *X. laevis* gene models (Fig. 8D; Table S4). Of them, 1,252 TF genes whose maximum TPM is 2 or more were subjected to HCA using the cutree function in *R.* In particular, we focused on 10 pylorus/sphincter-related TF genes as benchmarks, *barx1, cdx2, gata3, nkx2-3, nkx2-5, nkx3-2, pdx1, six2, sox2,* and *sox9*, in which *nkx2-3, pdx1, six2,* and *sox2* were added, according to the literature (Le Guen et al., 2015; Udager et al., 2010), but *gata3* was excluded due to its expression lower than 2 TPM. We observed how these 9 pylorus/sphincter-related TF genes were clustered with each other or with other genes by varying the number of clusters as a parameter, and found that 50 clusters were suitable for further analysis, as follows (Fig. 8E; Table S5). Under this condition, 6 out of 9 pylorus/sphincter-related TF genes, *nkx2-3.L/S, nkx2-5.L/S, nkx3-2.L/S, pdx1.L/S, six2.L/S,* and *cdx2.L*, were clustered into a single cluster, #2; *barx1.L, barx1.S, sox2.L/S,* and *sox9.L* were separately clustered into #8, #14, #42, and #39, respectively. Notably, L and S homeologs of *nkx2-3, nkx2-5, nkx3-2, pdx1,* and *six2* in #2 as well as *six2* in #42 were clustered together, consistent with the notion that expression profiles of L and S homeologs of developmentally regulatory TF genes in *X. laevis* are highly correlated with each other in more than 80% of 189 homeolog pairs (Watanabe et al., 2017). Therefore, we assumed that clusters with similar expression profiles and those containing homeolog counterparts could be regarded as related clusters. Based on this, clusters #2, #8, #14, #42, and #39 were assembled into three groups with other related clusters as follows (Fig. S7; Table S6). Group 1 (clusters #2 and #39) includes 7 of the pylorus/sphincter-related TFs (*nkx3-2.L/S, pdx1.L/S, nkx2-5.L/S, six2.L/S, nkx2-3.L/S, cdx2.L,* and *sox9.L*) (Figs. 8F, S8A-J), and show an ascending anterior-to-posterior (AP) gradient expression before and during metamorphosis (Fig. S7A,B). Group 2 (clusters #8, #14, #1, and #15) included *barx1.L/S* as well as *hand2.L/S, rxrb.L/S, snail2.L/S, sox4.L/S* (Figs. 8H, S8K) and are broadly upregulated in the A through PY regions at stage 60 (Figs. S7C-F, S8K-S). Group 3 (clusters #42, #6, #21, and #29) included *sox2.L/S* (Figs. 8J, S8U), and were characterized by an ascending AP gradient expression and downregulation during metamorphosis (Figs. 8J,K, S8U-AL). These groups are expected to include uncharacterized TF genes that could negatively or positively regulate pyloric sphincter formation during pre-metamorphic or metamorphic climax stages, respectively.

In Group 1, *nkx3-2.L/S* and *pdx1.L/S* in cluster #2 were highly expressed in the posterior stomach at both stages 53 and 60 (Figs. 8F, S7A, S8A,B), whereas *nkx2-5.L/S* was specifically expressed in the pylorus region (Fig. S8C), implying that cluster #2 TFs specify the pylorus region before and during metamorphosis. *pdx1.L* and *cdx2.L* were downregulated at stage 60 as st60_1 and _2 samples (closed dots) indicated but not st60_3 (early) (open triangles) (Fig. S8B,F, left panels). As mentioned above (see Fig. 8B), st60_3 (early) samples were likely to represent an earlier stage than st60_1 and _2 samples, suggesting that downregulation of *pdx1.L* and *cdx2.L* occurs relatively late in stage 60. Notably, cluster #2 included a reprogramming factor *klf4.L/S* (Bialkowska et al., 2017) and an AP patterning factor *meis2.L* (Schulte and Geerts, 2019) (Fig. 8G, S8H,I), whose roles in pylorus formation have not been investigated yet. Cluster #39 including *sox9.L* and *klf6.S* (a tissue remodeling factor; Bialkowska et al., 2017) showed a similar expression pattern with cluster #2 at stage 53, but was broadly upregulated in the stomach at stage 60 (Figs. S7B, S8G,J). These results suggest that Group 1 genes include positive regulators for sphincter formation.

Group 2 genes were characterized by a broad upregulation in every stomach region at stage 60 (Figs. S7C-F, S8K-T), but the mode and timing of upregulation were in several patterns. *thibz.L/S* and *sox4.L/S* were barely expressed at stage 53, but strongly induced at stage 60 (Fig. S8L,M), whereas *barx1.L/S* and other genes were relatively weakly expressed at stage 53, and upregulated at stage 60 (Fig. S8K). Regarding the timing, the upregulation of most genes in cluster #8, such as *barx1.L, thibz.L/S, sox4.L/S*, and *hand2.S*, appeared to occur relatively late in stage 60 because expression levels in st60_3 (early) samples (open triangles) were lower than those of the st60_1 and _2 samples (closed circles) in all A, M, P, and PY regions (Fig. S8K-N; see also heatmaps marked by open triangles in Fig. S7C-F). In fact, *thibz* (also known as *TH/bZip*) is reportedly a late response gene directly induced by TH in metamorphosis and has been proposed to promotes cell proliferation of the adult epithelial primordia of the stomach and intestine (Buchholz et al., 2005; Ishizuya-Oka et al., 1997). By contrast, upregulation of most genes in cluster #15, such as *rxrb.S, rara.L/S,* and *cebpd.L/S,* were likely to occur early in stage 60 as judged by the st60_3 (early) samples. Cluster #14 and #1 genes, such as *hand2.L, snai2.L/S*, *twist1.L/S*, and *foxf1.L/S*, had more divergent expression levels between st60_1 and _2 in addition to st60_3 (early) compared to #8 and #15 (Fig. S8N-Q); furthermore, the relative level of st60_3 (early) expression was not always lower than those of st60_1 and _2 samples (e.g., compare st60A with st60M in *hand2.L, snai2.L/S, twist.L/S* in Fig. S8N-P), possibly due to the difference in the timing of gene induction among the regions. Thus, it is likely that Group 2 TF genes include both early and late upregulated genes, which might be either directly or indirectly induced by TH during metamorphosis.

Group 3 genes were highly expressed in the posterior stomach and were downregulated during metamorphosis (Fig. S7G-J), as typically exemplified by *sox2.L/S, foxq1.S, foxa1.S, nkx6-2.S,* and *nkx6-3.S* (Figs. 8J,K, S8U-Y). In these genes, expression levels of the st60_3 (early) samples (open triangles) were higher than the other two (closed circles), suggesting their downregulation occurs late in stage 60. Since Foxq1 reportedly functions as a transcriptional repressor (Kang et al., 2019), downregulation of *foxq1.S* at stage 60 may release the suppression machinery of sphincter formation (Fig. 8K). *meis1.L* and *gbx1.L* were expressed in an ascending AP gradient manner at stage 53, but this gradient was diminished or lost at stage 60 (Fig. S8Z,AA). These data imply that Group 3 genes confer posterior identities of the pre-metamorphic stomach and negatively regulate sphincter formation.

We further examined whether the pre-metamorphic stomach expresses “intestinal TF genes” like *cdx2.L* (Figs. 4, 7B,C, S8F). We checked 24 mouse embryonic intestinal TF genes (including *cdx2*) identified from microarray and other analyses (Gao et al., 2009; Li et al., 2009); reviewed in Le Guen et al. (2015a); Table S8). Of them, 17 genes (not considering L and S genes) were expressed in the pre-metamorphic (stage 53) and metamorphic (stage 60) stomach at 2 TPM or higher, and were classified into three categories: upregulated, downregulated, and unchanged at stage 60 compared to stage 53 (Table S9, Fig. S8AB-AP). Upregulated genes included four Group 2 genes, *foxf1.L/S, foxf2.L/S, mafb.L/S,* and *tfec.S* (Fig. S8Q,AB-AD) as well as *hnf4a.L/S* and *nr3c2.L* (Fig. S8AE,AF), whereas downregulated genes included two Group 1 genes (*cdx2.L* and *hnf4g.L;* Fig S8F,AG), three Group 3 genes (*irf7.L, isx.L1,* and *mlxipl.L*; Fig. S8AH-AJ) as well as *srebf1.L* (Fig. S8AK), and unchanged genes included *nr1i3.S* (Group 3) and the other 6 genes (Fig. S8AL-AP). Of note, ten of the 17 intestinal genes belonged to Groups 1-3, implying their possible involvement in sphincter formation (Table S9). Expression analysis in adult *Xenopus* using public RNA-seq datasets (Session et al., 2016) showed that 9 genes, such as *cdx2.L, hnf4a.L/S*, *hnf4g.L/S*, *mafb.L/S,* and *srebf1.L/S,* were preferentially expressed in the intestine compared to the stomach (Table S10; also indicated in magenta in Table S9). These genes could be used as intestinal markers in the tadpole to adult in *Xenopus*. Our results showed that various intestinal genes are expressed in the posterior stomach of *Xenopus* before the stomach-duodenum boundary is formed. This may explain the transformation of the posterior stomach to the transitional zone in frogs (Barrington, 1946; Bodegas et al., 1997; Chalmers et al., 2000), that is possibly homologous to the intermediate zone before sphincter formation in mice (Li et al., 2009).

We next focused on cell-cell signaling pathways of Hedgehog (Hh) (34 genes), TGF-β superfamily (196 genes), Wnt (172 genes), FGF (111 genes), and RA (36 genes) (Michiue et al., 2017; Suzuki et al., 2017; Table S7). *Shh* and *bmp4* are sequentially induced by TH in the metamorphic intestine (Ishizuya-Oka et al., 2006; Stolow and Shi, 1995). Consistently, *shh.L*/*S* and *bmp4.L* were upregulated in this order at stage 60 (Fig. S9A,D). *shh.L/S* as well as *bmp4.L, bmp7.L, nog.S, bmp1.L/S, wnt5a.L/S,* and *fgf19.S* expression showed a descending AP gradient (Fig. S9A,D-F,I,J,M), whereas *ihh.L*, *bmpr1b.L/S, wnt5b.L, a2m.S* (*endodermin.S*)*, cyp26b1.L* expression showed an opposite gradient (Fig. S9B,H,K,O,Q). *wnt5b.S* was upregulated at stage 60 and showed high expression in M and P (Fig. S9K, right panel). *shh* and *ihh* expression forms a reciprocal gradient (Fig. S9A,B), which is also reported in the mouse stomach (Kolterud et al., 2009). For RA pathways, *aldh1a1.L*, an RA-synthesizing enzyme, was strongly expressed in the posterior stomach at stage 53, but was downregulated at stage 60 (Fig. 8L; the same as Fig. S9P, left panel). By contrast, *cyp26b1.L*, an RA-degrading enzyme, was upregulated in the posterior stomach at stage 60 (Fig. 8M; st60M, P, and PY; the same as Fig. S9Q, left panel), suggesting that the amount of RA is reduced in the posterior stomach during metamorphosis. Furthermore, nuclear RA receptor genes, *rara.L/S* and *rxrb.L/S*, whose products form the RAR/RXR heterodimer, were broadly expressed in the stomach and were upregulated at stage 60 (Fig. S8R,S). Altogether, these data showed that *shh.L*/*S*, *ihh.L*, *bmp4.L*, *bmp7.1.L*, *bmpr1b.L/S*, *chrdl1.L*, *wnt5b.L, fgf19.S*, *fgfr2*, *a2m.S,* and *cyp26b1.L* may promote sphincter formation, while *dhh.L/S*, *aldh1a1.L*, *rara.L/S* and *rxrb.L/S* may inhibit it.

In summary, through the spatiotemporal transcriptome analysis, we identified uncharacterized genes for various TFs and cell-cell signaling factors, which could provide insight into the molecular mechanism of how pyloric sphincter formation is suppressed in embryogenesis in herbivorous tadpoles but promoted during metamorphosis in carnivorous adults. Of the particular interest is RA signaling, which might be a key factor in suppressing sphincter formation, as discussed below.

## Discussion

In this paper, histological and molecular analyses using *X. laevis* revealed the timing and place of sphincter formation in the pylorus region (Fig. 9A). Furthermore, the region-specific RNA-seq analysis on the pre-metamorphic and metamorphic stomach led to the identification of previously uncharacterized genes that may be involved in the formation of the sphincter and pylorus-duodenum boundary. Based on these findings, we discuss the morphological and molecular mechanisms of the pylorus development in anurans (Fig. 9B).

**Fig. 9.**
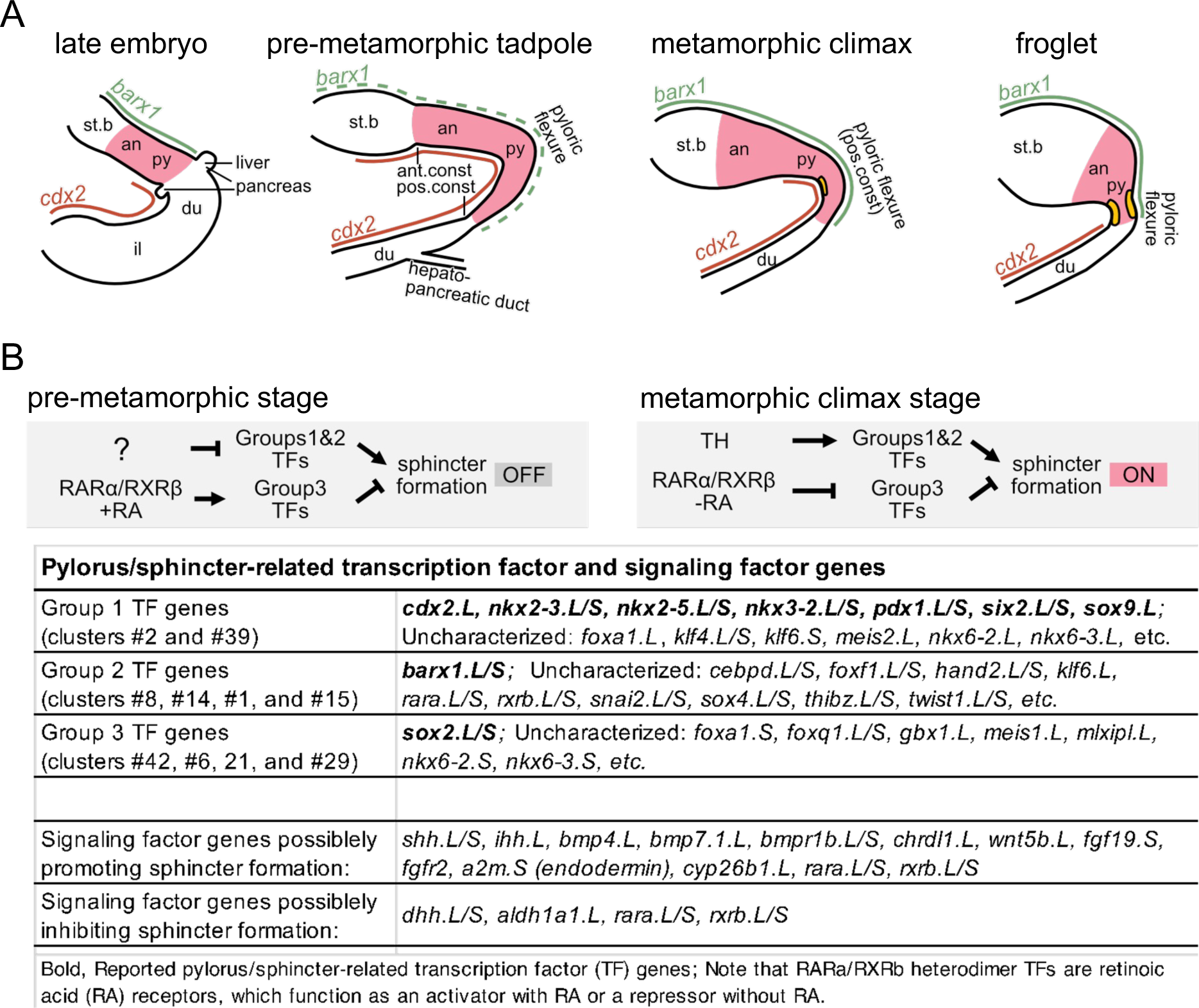
Morphogenesis of the stomach-duodenum region in *Xenopus* development and a possible molecular mechanism of pyloric sphincter formation during metamorphosis. (A) Morphological and molecular features of the stomach-duodenum region during development. Late embryos (stage42/43), pre-metamorphic tadpoles (stages 52-55), metamorphic climax (stages 59-66) and froglet (st.66). Elongation and shortening of the posterior stomach (antrum and pylorus region) is highlighted in pink. Yellow bands, the pyloric sphincter. an, antrum; ant.const, anterior constriction; du; duodenum; st.b, stomach body; il, ileum; pos.const, posterior constriction; py, pylorus region. (B) A hypothesis for the molecular mechanism of pyloric sphincter formation during metamorphosis. Arrows, activation; T marks, inhibition. Groups 1, 2, or 3 transcription factor (TF) and retinoic acid (RA) signaling-related genes are identified by RNA-seq analysis (see Figs 8, S8, S9), which include known pylorus/sphincter-related genes as well as uncharacterized genes possibly involved in sphincter formation. Representative TF genes in Groups 1, 2, or 3 and signaling factor genes are shown in the table below. See the text for details.

### Formation of the pyloric sphincter during *Xenopus* metamorphosis

Our detailed histological analyses revealed that the pyloric sphincter starts to be formed at stage 61 or slightly earlier on the inner (dorsal) side of the flexure within the posterior stomach, and progresses to the outer (ventral) side as metamorphosis proceeds (Figs. 2D, 9A). This flexure is often called the gastroduodenal (GD) loop, which was thought to compartmentalize the stomach and intestine (Bloom et al., 2013; Womble et al., 2016). However, judging from the expression patterns of the stomach marker *barx1* and the pylorus-related genes *sox9*, *nkx3-2*, *gata3*, *grem1*, and *nkx2-5*, the flexure clearly belongs to the pylorus region at least from stage 42/43 onward (Figs. 4-6), so hereafter it will be called the pyloric flexure. The progression of pyloric sphincter formation from the inner curve (dorsal) toward the outer curve (ventral) side of the pylorus is a new finding in this study. Considering the presence of the typhlosole-like structure in the inner curve side of the pyloric flexure (Ishizuya-Oka et al., 1997), thickening of the circular muscle for sphincter formation may start at the typhlosole-like structure. This speculation comes from the observation that the thickened muscle is surrounded by a connective cell mass on the dorsal side (Fig. 2D,E1), but not on the ventral side (Fig. 2E2).

### Morphogenesis of the pylorus-duodenum region in *Xenopus*

The pyloric and duodenal epithelia are morphologically similar at embryonic and tadpole stages (Figs. 1, 2), but after metamorphosis they are clearly distinguished and separated by the sphincter (Fig. 3). The molecular boundary is also unclear at embryonic and tadpole stages, because expression of *barx1* and the intestinal genes such as *cdx2*, *hnf4g*, *maf*, *mafb*, and *prrx1* overlap in the pylorus region (Figs. 4, 7, S8F,AB-AP; Table S9), like the intermediate zone of the mouse posterior stomach, where many intestinal genes are expressed before E16.5 (i.e. before pyloric sphincter formation; Li et al., 2009). Therefore, formation of the stomach-intestine boundary at morphological and molecular levels may require suppression of intestinal genes in the posterior stomach. In mice, Barx1 suppresses *Cdx2* expression in the stomach through *Sfrp1* and *Sfrp2* induction (Kim et al., 2005). The same regulatory system may be activated in the *Xenopus* posterior stomach during metamorphosis, while it is suppressed before metamorphosis possibly by TFs that are downregulated during metamorphosis (Figs. 8, S8). Of note, suppression of *cdx2* expression in the stomach and the epithelial pylorus-duodenum boundary formation occurs after the onset of pyloric sphincter formation in both mice (Li et al., 2009; Udager et al., 2010) and *Xenopus* (this study), so the formation of the molecular boundary of the pylorus-duodenum region may not be a trigger for sphincter formation, but is independent or downstream of it.

We further revealed that the pylorus region in *Xenopus* undergoes dynamic remodeling by elongating at embryonic to early tadpole stages and shortening during metamorphosis (Fig. S1), as has been shown for the intestine (Schreiber et al., 2005). The elongation of the GI tract of tadpoles is regarded as an adaptation for herbivory. In changing to carnivory, the pyloric sphincter starts to be formed at the far end of the inner curvature of the pyloric flexure within the pylorus region during metamorphosis, but is eventually located at the posterior end of the pylorus by the end of metamorphosis through the shortening between the pyloric flexure and posterior constriction (Fig. 9A). Thus, the pylorus region shares various features with the duodenum, such as similar epithelial morphology (Fig. 1), intestinal gene expression (Figs. 4, 7; Table S9), and the shortening during metamorphosis (Figs 2, 3).

### Mechanism of the heterochrony of pyloric sphincter formation

A remarkable characteristic of the pyloric sphincter in anurans in contrast to amniotes is that the timing of sphincter formation is temporally separated from embryonic morphogenesis and suspended until metamorphosis. There are two possible mechanisms underlying this heterochrony: (i) essential genes for sphincter formation are not expressed before metamorphosis, and (ii) repressive genes against sphincter formation are expressed before metamorphosis. Most of the pylorus/sphincter-related genes in amniotes such as *barx1, nkx3-2, nkx2-5, sox9, gata3, and grem1* are already expressed before metamorphosis (Figs. 4, 5, 9B), and therefore do not fit the criteria for mechanism (i). Instead, we have newly identified TFs that are upregulated in the metamorphic pylorus region, such as *ascl1.*L/S, *foxa1.L*, *klf3.L*, *klf4.L/S*, *nkx6-2.L*, *nkx6-3.L*, *pax6.L*, *rara.L/S*, and *rxrb.L/S*, *sox4.L/S*, *thibz.L/S*, all of which belong to Group 2 (Figs. 9B, S7, S8). Of them, *nkx6-2* is reportedly expressed in the embryonic stomach in chick and *Xenopus* (Dichmann and Harland, 2011; Pedersen et al., 2005), and hence may not have an initiative role in sphincter formation. TF genes in this list that are not expressed in the embryonic stomach will be good candidates for mechanism (i). Regarding mechanism (ii), repressor TFs genes in Group 3, such as *foxq1* as mentioned above (Kang et al., 2019; (Fig. 8K), could be involved. Identifying those TF genes will help reveal how sphincter formation is initiated during *Xenopus* metamorphosis.

A promising candidate involved in mechanisms (i) and (ii) is RA signaling, because it was reported that inhibition of RA signaling transforms the stomach of the *Xenopus* larva into a stomach resembling the carnivorous *Lepidobatrachus laevis* larva, which possesses a pyloric sphincter (Bloom et al., 2013; Ruibal and Thomas, 1988). This raises the possibility that RA signaling suppresses sphincter formation, and its reduction triggers sphincter formation. Consistent with this hypothesis, the RA-synthesizing and degrading enzyme genes, *aldh1a1.L* and *cyp26b1.L*, are up- and down-regulated, respectively, (leading to a high RA level) in the posterior stomach before metamorphosis, whereas those genes are down- and up-regulated (leading to a low RA level) during metamorphosis (Fig. 8L, M). The RA receptor is a heterodimer of RARα and RXRβ and functions as a transcriptional activator or repressor depending on the presence or absence of RA, respectively (Hauksdottir et al., 2003). Notably, the receptor genes, *rara.L/S* and *rxrb.L/S*, are expressed before metamorphosis and further upregulated during metamorphosis (Fig. S8R,S). It is therefore tempting to speculate that RARα/RXRβ heterodimers act as a molecular switch by activating the suppressive machinery for sphincter formation before metamorphosis when RA is abundant and inhibiting it during metamorphosis once RA is reduced (Fig. 9B). This hypothesis could be tested by examining whether sphincter formation is initiated prematurely by adding RA inhibitors before metamorphosis or is inhibited by RA treatment during metamorphosis.

In conclusion, our study provided detailed histological and molecular descriptions of the pylorus region and sphincter development in *Xenopus*, constructed RNA-seq datasets that serve as a useful resource for future studies on gastric remodeling, and proposed hypotheses on how the sphincter is formed during anuran metamorphosis. Testing these hypotheses would also provide insights into the question of how the heterochronic changes in the timing of sphincter formation were achieved during the evolutionary diversification of vertebrate GI tracts.

## Data access

All sequencing data generated in this study have been submitted to the DDBJ Sequence Read Archive (DRA) database (https://www.ddbj.nig.ac.jp/dra/index.html) under accession number DRA016712.

## Acknowledgements

We thank Katsushi Yamaguchi for supporting RNA-seq experiments; Takayoshi Yamamoto for pCSf107-BMP4S-T; Jean-Pierre Saint-Jeannet for pGEMT-Xsox9; Mariko Kondo for supporting RNA-seq experiments and critical reading of the manuscript. We also thank the National BioResource Project (NBRP) of MEXT for the database of *Xenopus laevis* and *Xenopus tropicalis*. This work was supported by the Sasakawa Scientific Research Grant from The Japan Science Society (K.N.), JSPS KAKENHI under Grant Numbers JP 25251026 (M.T.), and NIBB Collaborative Research Program 18-414 (M.T.).

## Author contributions

Conceptualization, K.N. and M.T.; Methodology, K.N., T.I., T.H., Y.S-K. S.S. and M.T.; Investigation, K.N., T.I., S.U., and M.T.; Resources, K.N.; Writing - Original Draft, K.N., T.I. and M.T.; Writing - Review & Editing, All authors; Supervision, A.I-O. and M.T.; Funding Acquisition, K.N. and M.T..

## Supplementary Information

### Supplementary Tables in Excel files

Table S1. Primer sequences for cDNA PCR cloning.

Table S2. Differentially expressed genes between stomach regions A, M, and P at stages 53 and 60.

Table S3. Gene ontology term annotation by EggNOG mapper v2.

Table S4. List of transcription factor (TF) genes in *Xenopus laevis*.

Table S5. RNA-seq expression profiles of TF genes in *Xenopus laevis*.

Table S6. Representative genes and L/S pairs in Groups 1, 2, and 3 associated with pylorus/sphincter-related TF genes.

Table S7. RNA-seq expression profiles of signaling factor genes in *Xenopus laevis*.

Table S8. List of amniote embryonic intestinal TF genes and *Xenopus* orthologs.

Table S9. Expression profiles of intestinal TF genes in pre-metamorphic and metamorphic stomach.

Table S10. Differentially expressed gene analysis of embryonic intestinal TF genes in adult *Xenopus* intestine and stomach.

### Supplementary Figure legends

Fig. S1. External morphology of the gastrointestinal tract and the stomach-duodenum region during *Xenopus* development.

External morphology of the gastrointestinal (GI) tract (A-C, F) and the stomach-duodenum region (D, E) at stage 42/43 (A), stage 46 (B, C), stage 53 (D), stage 60 (E), and stage 63 (F) in *Xenopus laevis*. The GI tract at stage 63 (F) is yellowish due to fixation by Bouin’s solution. Magenta asterisk, flexure or pyloric flexure (see Discussion). Magenta arrow, posterior constriction. Magenta arrowhead, anterior constriction; white arrowhead, position of the bile duct opening. an, antrum; p.st, posterior stomach; du, duodenum; il, ileum; l.b, liver bud; p.b, pancreatic bud, py, pylorus region; st, stomach; st.b, stomach body. Scale bars, 1 mm.

Fig. S2. Histology of the pylorus region at metamorphic climax (stage 61).

(A) Low-magnification image of Fig. 2D. Boxes 1, 2, and 3 indicate positions of Fig. 2D1-3. (B) A wider view of Fig. 2D2. Anterior to the left. Magenta asterisk, pyloric flexure. c.t, connective tissue; ep, epithelium. Scale bars, 100 μm.

Fig. S3. *acta2* as a positive control for section in situ hybridization.

*acta2* anti-sense DIG probe was hybridized with a section from the same serial sections as *barx1* in situ hybridization around the anterior constriction at the pre-metamorphic tadpole (stage 53). Magenta arrowhead, anterior constriction. an, antrum; st.b, stomach body. Scale bar, 100 μm.

Fig. S4. Dissection of gastrointestinal tracts for RT-PCR analysis at pre-metamorphic and metamorphic climax stages.

Gastrointestinal (GI) tracts dissected at pre-metamorphic tadpole stage 53 (A) and metamorphic climax stage 60 (B) for RT-PCR analysis (Fig. 7A) are shown. Magenta asterisk, pyloric flexure. Magenta arrow, posterior constriction. Magenta arrowhead, anterior constriction. b.d, bile duct. Scale bars, 1 mm.

Fig. S5. Hierarchical clustering and gene ontology analyses of RNA-seq data from dissected stomach regions at pre-metamorphic and metamorphic climax stages.

RNA-seq data obtained from dissected stomach samples, A (anterior), M (mid), and P (posterior) parts (see Fig. 8A), at pre-metamorphic stages 53 (st53) and metamorphic climax 60 (st60) were used for hierarchical clustering analysis (HCA) and gene ontology (GO) analysis. The sample name st53A_1, for example, indicates stage 53, anterior part, replicate #1. (A) HCA. Euclidean distances between samples are shown as color-coded heatmaps. Note that st60_3 (early) data (dark blue box) are clustered with st53 data rather than st60_1 and st60_2. Euclidean distances between st53A_1_2_3 and st53P_1_2_3 transcriptomes (pale blue box) and those between st60A_1_2 and st60B_1_2 (green box) were plotted in Fig. 8C. (B, C) GO analysis. Lolliplots show fold enrichment of GO terms in genes that are significantly upregulated (B) or downregulated (C) in st60P samples (including st60P_3) compared with st53P samples. X-axis, fold enrichment; the color of the circle, −log10 (adjusted p-value); the size of the circle, the number of genes. The top 20 terms ordered by fold enrichment values are shown.

Fig. S6. Differentially expressed gene analysis of RNA-seq data between pre-metamorphic and metamorphic climax stages.

(A-I) Volcano plots of differentially expressed genes (DEGs) between anterior (A), mid (M), and posterior (P) regions of the stomach at pre-metamorphic (stage 53) and metamorphic climax (stage 60) stages are shown. The numbers of up- and down-regulated DEGs (e.g. A vs M means up- and down-regulation in M compared to A) between two regions or stages are indicated by the numbers with ‘up’ or ‘down’ in the plots. X-axis, log2 fold change; y-axis, −log10 (padj). Red dots, DEGs. The top 10 DEGs from the highest −log10 (padj) value are labeled with gene names. (A-F) Comparison between regions A, M, and P at stage 53 (A-C) and at stage 60 (D-F). Note that the numbers of up- and down-regulated DEGs are both much smaller at stage 60 than at stage 53 (Compare panels A and D, panels B and E, and panels C and F; see also Table S2). The total number of genes plotted are indicated in each panel. This may reflect that regional differences are equalized by dedifferentiation and remodeling of the entire stomach during metamorphosis (see the text). (G-I) Comparison of regions A, M, or P between stages 53 and 60. The numbers of both up- and down-regulated DEGs of stages 53 vs 60 in each region are larger than those between regions at stage 53 or 60 (A-F). Noticeable DEGs at stage 60 are *sox4.L/S*, *thibz.S*, and *shh.L* as up-regulated genes (underlined in G,H,I) and *atp4b.L/S* as down-regulated genes (magenta box in E,F,H). *Sox4.L/S, thibz.S,* and *shh.L/S* are reportedly upregulated by thyroid hormone (TH) in the entire stomach at stage 60 (see also Figs. S8M,N, S9A). (J, K) Dot plots of *atp4a.L/S* (J) and *atp4b.L/S* (K). *atp4a* and *atp4b* encode gastric H(+)-K(+)-ATPase subunits alpha and beta, respectively, which form a proton pump to generate acidic gastric juice in the stomach body; therefore, *atp4a* and *atp4b* genes are used as terminal differentiation markers for the stomach body (Willet and Mills, 2016). Down-regulation of *atp4a.L/S* and *atp4b.L/S* indicates the elimination of differentiated cells and/or the transition to a dedifferentiation state and is an example of the equalization of gene expression profiles in the stomach during metamorphosis. Y-axis, transcripts per million (TPM).

Fig. S7. Selected clusters of transcription factor genes identified by hierarchical clustering analysis of regional expression profiles in the stomach at pre-metamorphic and metamorphic climax stages.

Transcription factors (TF) genes with 2 or more TPM (1,252 out of 2,435 genes) were divided into 50 clusters by hierarchical clustering analysis (HCA) (see Fig. 8E; Table S5), and 10 out of them were assembled into three groups (Tables S5, S6; see the main text for details). (A,B) Group 1 gene clusters, #2 and #39. (C-F) Group 2 gene clusters, #8, #14, #1, and #15. (G-J) Group 3 gene clusters, #42, #6, #21, and #29. Representative genes are shown in the parentheses. The pylorus/sphincter-related genes are indicated in magenta and by magenta boxes. Note that too many genes are not all listed on the right side of the heatmaps (C,E,F). See Table S5 for all clusters and their consisting genes. Expression level is shown by color-coded z-scored TPM. Vertical dotted line separates pre-metamorphic stage 53 and metamorphic climax stage 60 samples.

Fig. S8. Dot plot analysis of regional and temporal expression profiles of transcription factor genes in the stomach at pre-metamorphic and metamorphic climax stages.

Dot plots of expression profiles are shown for representative TF genes from the selected 10 clusters (see Fig. S7) and from intestinal TF genes (see Table S9). The ordinate indicates transcripts per million (TPM). The sample names, st53A, st53M, st53P, st53PY, st60A, st60M, st60P, st60PY, are indicated below the plot. (A-J) Group 1 genes (clusters #2 and #39). (K-T) Group 2 genes (clusters #8, #14, #1, and #15). (U-AA) Group 3 genes (clusters #42, #6, #21, and #29). (AB-AP) Intestinal genes. Those are subdivided into upregulation (AB-AF), downregulation (AG-AK), and no change (AL-AP). (A) *nkx3-2,* (B) *pdx1,* (C) *nkx2-5,* (D) *six2,* (E) *nkx2-3,* (F) *cdx2,* (G) *sox9,* (H) *klf4,* (I) *meis2*, (J) *klf6,* (K) *barx1,* (L) *thibz (thyroid hormone induced bZip protein),* (M) *sox4,* (N) *hand2,* (O) *snai2,* (P) *twist1*, (Q) *foxf1,* (R) *rxrb (retinoid X receptor beta),* (S) *rara (retinoic acid receptor alpha),* (T) *cebpd* (*CCAAT enhancer binding protein delta*), (U) *sox2,* (V) *foxq1,* (W) *foxa1,* (X) *nkx6-2,* (Y) *nkx6-3,* (Z) *meis1*, (AA) *gbx1*, (AB) *foxf2*, (AC) *mafb*, (AD) *tfec* (*transcription factor EC*), (AE) *hnf4a* (*hepatocyte nuclear factor 4 alpha*), (AF) *nr3C2* (*nuclear receptor subfamily 3 group C member 2*), (AG) *hnf4g* (*hepatocyte nuclear factor 4 gamma*), (AH) *irf7* (*interferon regulatory factor 7)*), (AI) *isx* (*intestine-specific homeobox*), (AJ) *mlxipl* (*MLX interacting protein like*), (AK) *srebf1* (*sterol regulatory element binding transcription factor 1*), (AL) *hnf1a* (*HNF1 homeobox A*), (AM) *mlx* (*MAX dimerization protein MLX*), (AN) *nr1i3* (*nuclear receptor subfamily 1 group I member 3*), (AO) *ppara* (*peroxisome proliferator activated receptor alpha*), (AP) *prdm16* (*PR domain 16*). *sox2.S* exists as a fragment sequence LOC108718087 (gene37454) in the XENLA_9.2 gene model, although the full-length sequence was already cloned (Watanabe et al., 2017). Left and right plots in each panel are L and S homeologs in *Xenopus laevis*. Each plot is labeled with a group number followed by a cluster number (e.g. G1#2), only a cluster number (e.g. cluster #17) when it is outside of the Groups, or “<2TPM” when it is excluded from clustering. Mean TPM values are shown with magenta dots. Open triangles, st60_3 (early) samples; dotted lines in some panels, average values of upregulation (magenta) or downregulation (blue) without st60_3 (early) samples. In some cases, L and S genes are separately included in the two of the 10 clusters (e.g. *sox4, nkx6-2, nkx6-3, rxrb*) or only either L or S gene is included in one of the 10 clusters (e.g. *cdx2*, *sox9*) because they might have been subfunctionalized or their expression levels might have just fluctuated. Variations of three st60 points were much larger in certain genes (e.g. *barx1.L* in panel K and *thibz.L* in panel L) than the others (e.g. *rara.L/S* in panel R). In such cases, TPM values of st60_3 (early) (open triangles) were often between st53 samples and the other st60 samples, which is consistent with the suggestion that the actual stage of st60_3 is earlier than st60_1 and st60_2 (see the main text and Fig. 8B). The st60_3 (early) values can be used to infer the timing and/or velocity of up- or down regulation of genes during metamorphosis. For example, in *barx1.L* (K), TPM values of st60_3 (early) at all regions were lower than the other st60 points and closer to st53 points, suggesting that *barx1* upregulation is initiated after stage 53 and continues through stage60. By contrast, *rara.L/S* (S) is fully upregulated in the stage 60_3 (early) samples, suggesting that *rara.L/S* is induced in an early metamorphic climax stage. Expression levels of *pdx1.L* and *cdx2.L* in st60_3 (early) samples were all higher than st60_1 and 2 and rather similar to st53 samples, indicating that downregulation of these gene occurs relatively late in stage 60.

Fig. S9. Dot plot analysis of regional and temporal expression profiles of signaling factor genes in the stomach at pre-metamorphic and metamorphic climax stages.

Dot plots of expression profiles are shown for selected signaling factor genes. The ordinate, transcripts per million (TPM). The sample names, st53A, st53M, st53P, st53PY, st60A, st60M, st60P, st60PY are indicated below the plot. (A-C) Hh, (D-I) BMP, (J-L) Wnt, (M-O) FGF, (P, Q) Retinoic acid (RA) related genes. (A) *shh*, (B) *ihh*, (C) *dhh*, (D) *bmp4*, (E) *bmp7-1*, (F) *nog*, (G) *chrdl1*, (H) *bmpr1b*, (I) *bmp1*, (J) *wnt5a*, (K) *wnt5b*, (L) *tdgf1-2*, (M) *fgf19*, (N) *fgfr2*, (O) *a2m* (*alpha-2-macroglobulin*; the same as *endodermin*), (P) *aldh1a1*, (Q) *cyp26b1.* All ligand genes for Hedgehog, TGF-β superfamily, Wnt, and FGF signaling that are expressed at a maximum of 2 TPM or higher are shown (A-E, J, K, M). Left and right plots in each panel are L and S homeologs in *Xenopus laevis*. Mean TPM values are shown with magenta dots. Open triangles, st60_3 samples; dotted lines in some panels, average values of upregulation (magenta) or downregulation (blue) without st60_3 samples. For example, *bmp4.L, bmp7.L, chrdl1.L, wnt5b.S, fgf19.S, a2m.S* (D, E, K, M, O), st60_sample 3 (open triangle) showed lower values than st60_samples 1 and 2 at all positions, suggesting that their upregulation occurs at late stage 60. By contrast, *shh.L* and ihh.L (A,B), for example, st60_sample 3 showed values similar to st60_samples 1 and 2, suggesting that their upregulation occurs at early stage 60.

