## Supplementary material for "Histological and gene-expression analyses of pyloric sphincter formation during stomach metamorphosis in *Xenopus laevis*": Supplmentary Figures

### Supplementary information



Supplementary Tables (see each Excel file):

Table S1. Primer sequences for cDNA PCR cloning.

Table S2. Differentially expressed genes between stomach regions A, M, and P at stages 53 and 60.

Table S3. Gene ontology term annotation by EggNOG mapper v2.

Table S4. List of transcription factor genes in *Xenopus laevis.*

Table S5. RNA-seq expression profiles of transcription factor genes in *Xenopus laevis*.

Table S6. Representative genes and L/S pairs in Groups 1, 2, and 3 associated with pylorus/sphincter-related TF genes.

Table S7. RNA-seq expression profiles of signaling factor genes in *Xenopus laevis*.

Table S8. List of amniote embryonic intestinal genes and *Xenopus* orthologs.

Table S9. Expression profiles of intestinal genes in pre-metamorphic and metamorphic stomach.

Table S10. Differentially expressed gene analysis of embryonic intestinal genes in adult *Xenopus* intestine and stomach.

Supplementary Figures:

Fig. S1. External morphology of the gastrointestinal tract and the stomach-duodenum region during *Xenopus* development.

Fig. S2. Histology of the pylorus region at metamorphic climax (stage 61).

Fig. S3. *acta2* as a positive control for section in situ hybridization.

Fig. S4. Dissection of gastrointestinal tracts for RT-PCR analysis at pre-metamorphic and metamorphic climax stages.

Fig. S5. Hierarchical clustering and gene ontology analyses of RNA-seq data from dissected stomach regions at pre-metamorphic and metamorphic climax stages.

Fig. S6. Differentially expressed gene analysis of RNA-seq data between pre-metamorphic and metamorphic climax stages.

Fig. S7. Selected clusters of transcription factor genes identified by hierarchical clustering analysis of regional expression profiles in the stomach at pre-metamorphic and metamorphic climax stages.

Fig. S8. Dot plot analysis of regional and temporal expression profiles of transcription factor genes in the stomach at pre-metamorphic and metamorphic stages.

Fig. S9. Dot plot analysis of regional and temporal expression profiles of signaling factor genes in the stomach at pre-metamorphic and metamorphic stages.

Fig. S1 (Nagura et al.)

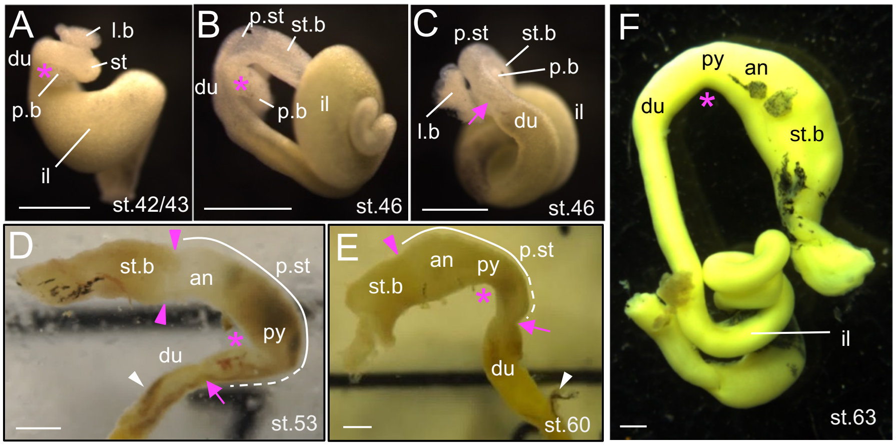


Fig. S1. External morphology of the gastrointestinal tract and the stomach-duodenum region during *Xenopus* development. External morphology of the gastrointestinal (GI) tract (A-C, F) and the stomach-duodenum region (D, E) at stage 42/43 (A), stage 46 (B, C), stage 53 (D), stage 60 (E), and stage 63 (F) in *Xenopus laevis*. The GI tract at stage 63 (F) is yellowish due to fixation by Bouin's solution. Magenta asterisk, flexure or pyloric flexure (see Discussion). Magenta arrow, posterior constriction. Magenta arrowhead, anterior constriction; white arrowhead, position of the bile duct opening. an, antrum; p.st, posterior stomach; du; duodenum; il, ileum; l.b, liver bud; p.b, pancreatic bud, py, pylorus region; st, stomach; st.b, stomach body; . Scale bars, 1 mm.

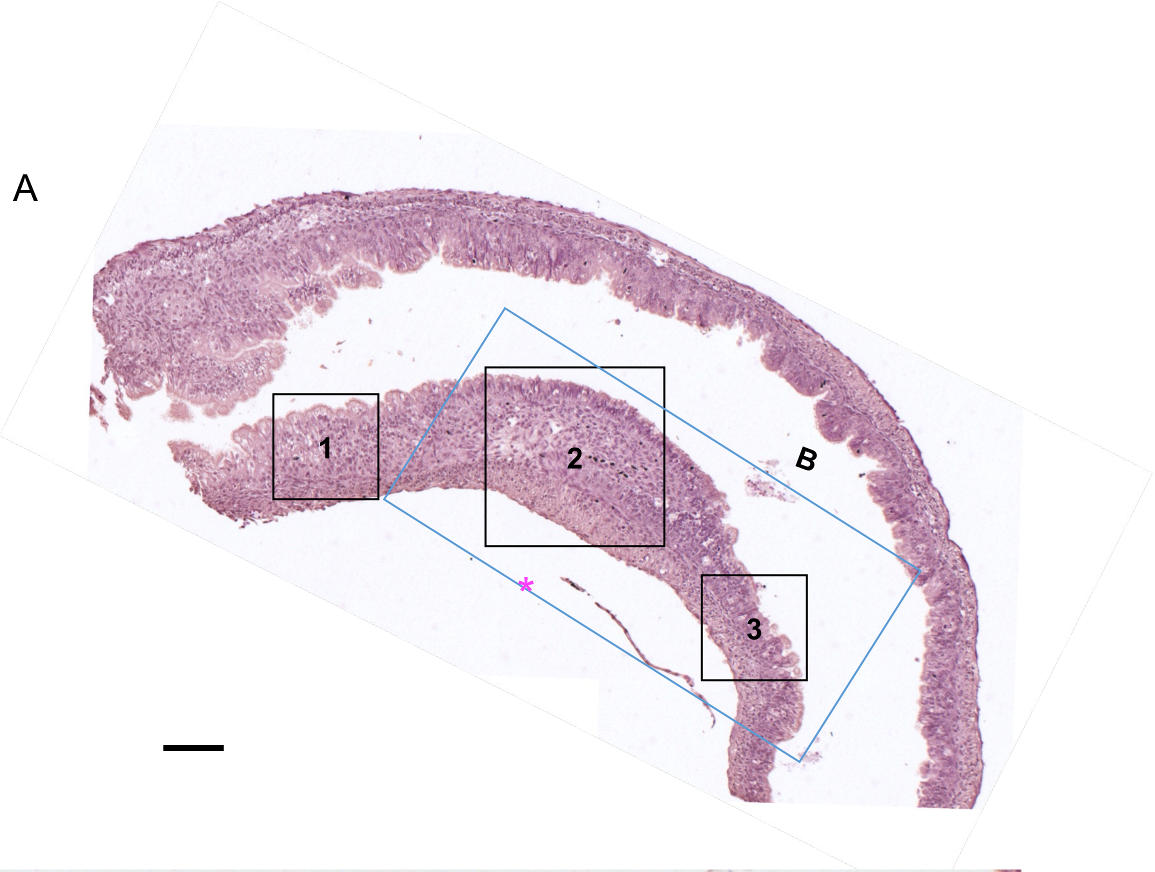


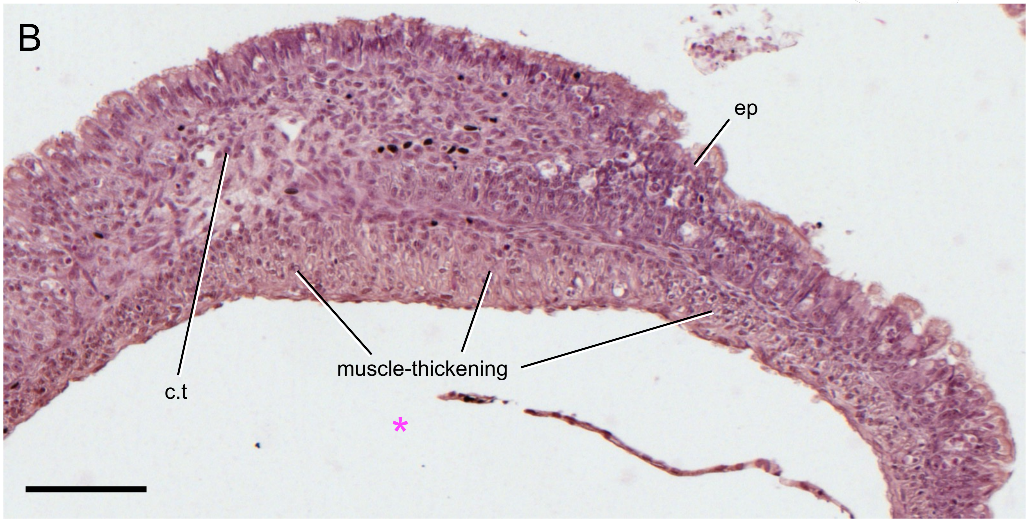


**Fig. S2. Histology of the pylorus region at metamorphic climax (stage 61).**

(A) Low-magnification image of Fig. 2D. Boxes 1, 2, and 3 indicate positions of Fig. 2D1-3. (B) A wider view of Fig. 2D2. Anterior to the left. Magenta asterisk, pyloric flexure. c.t, connective tissue; ep, epithelium. Scale bars, 100 µm.

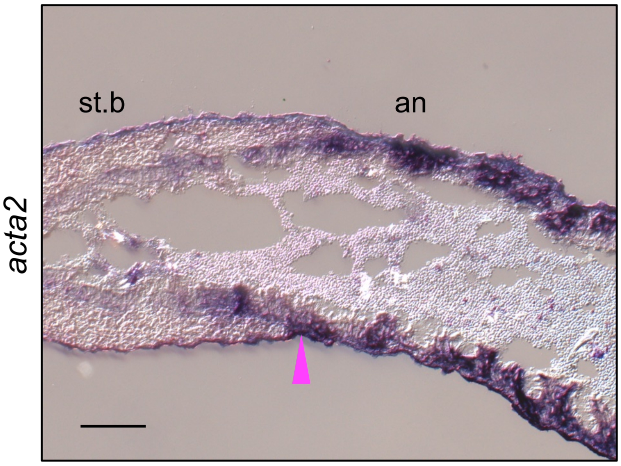


**Fig. S3. *acta2* as a positive control for section in situ hybridization.**

*acta2* anti-sense DIG probe was hybridized with a section from the same serial sections as *barx1* in situ hybridization around the anterior constriction at the pre-metamorphic tadpole (stage 53). Magenta arrowhead, anterior constriction. an, antrum; st.b, stomach body. Scale bar, 100 μm.

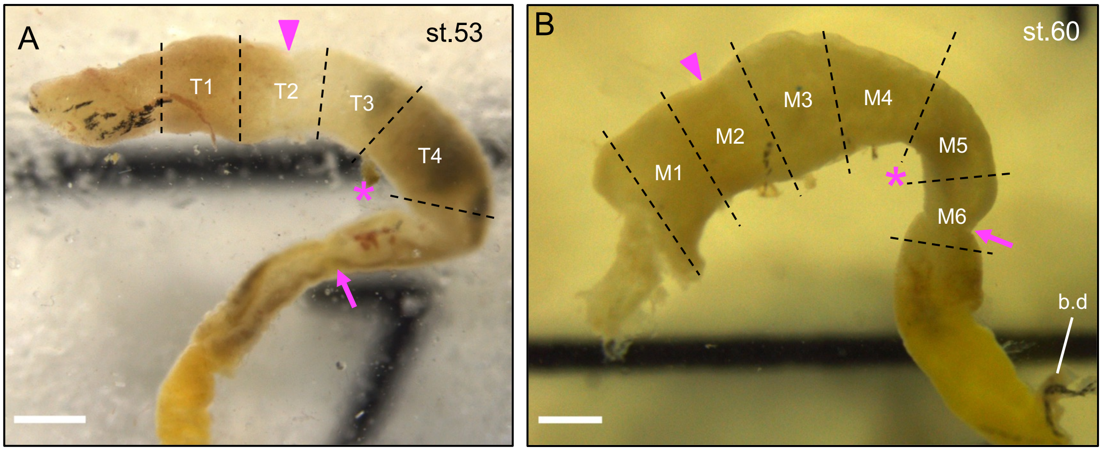


**Fig. S4. Dissection of gastrointestinal tracts for RT-PCR analysis at pre-metamorphic and metamorphic climax stages.**
Gastrointestinal (GI) tracts dissected at pre-metamorphic tadpole stage 53 (A) and metamorphic climax stage 60 (B) for RT-PCR analysis (Fig. 7A) are shown. Magenta asterisk, pyloric flexure. Magenta arrow, posterior constriction. Magenta arrowhead, anterior constriction. b.d, bile duct. Scale bars, 1 mm.

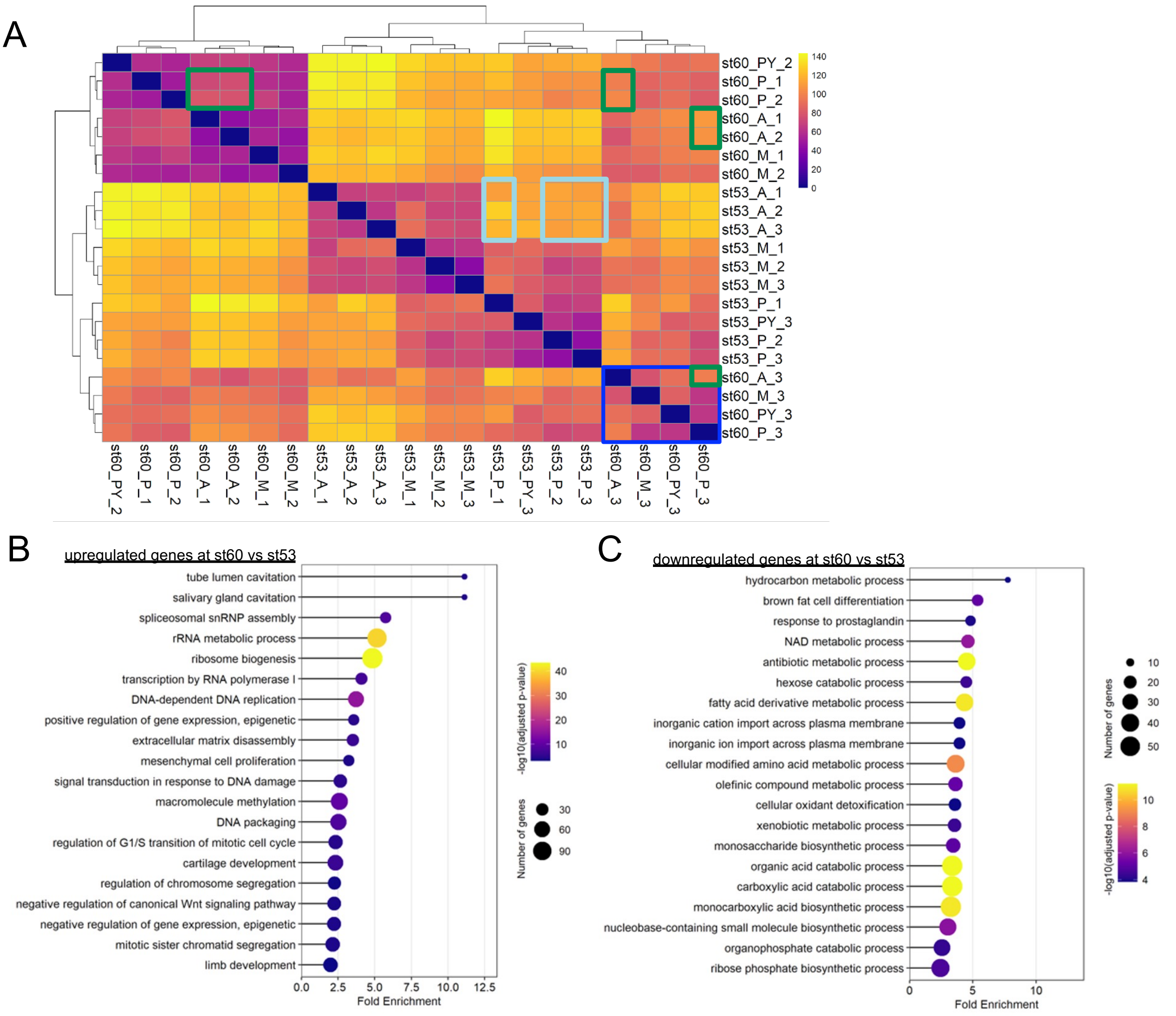


**Fig. S5. Hierarchical clustering and gene ontology analyses of RNA-seq data from dissected stomach regions at pre-metamorphic and metamorphic climax stages.**

RNA-seq data obtained from dissected stomach samples, A (anterior), M (mid), and P (posterior) parts (see Fig. 8A), at pre-metamorphic stages 53 (st53) and metamorphic climax 60 (st60) were used for hierarchical clustering analysis (HCA) and gene ontology (GO) analysis. The sample name st53A_1, for example, indicates stage 53, anterior part, replicate #1. (A) HCA. Euclidean distances between samples are shown as color-coded heatmaps. Note that st60_3 (early) data (dark blue box) are clustered with st53 data rather than st60_1 and st60_2. Euclidean distances between st53A_1_2_3 and st53P_1_2_3 transcriptomes (pale blue box) and those between st60A_1_2 and st60B_1_2 (green box) were plotted in Fig. 8C. (B, C) GO analysis. Lolliplots show fold enrichment of GO terms in genes that are significantly upregulated (B) or downregulated (C) in st60P samples (including st60P_3) compared with st53P samples. X-axis, fold enrichment; the color of the circle, -log10 (adjusted p-value); the size of the circle, the number of genes. The top 20 terms ordered by fold enrichment values are shown.

Fig. S6 (Nagura et al.)

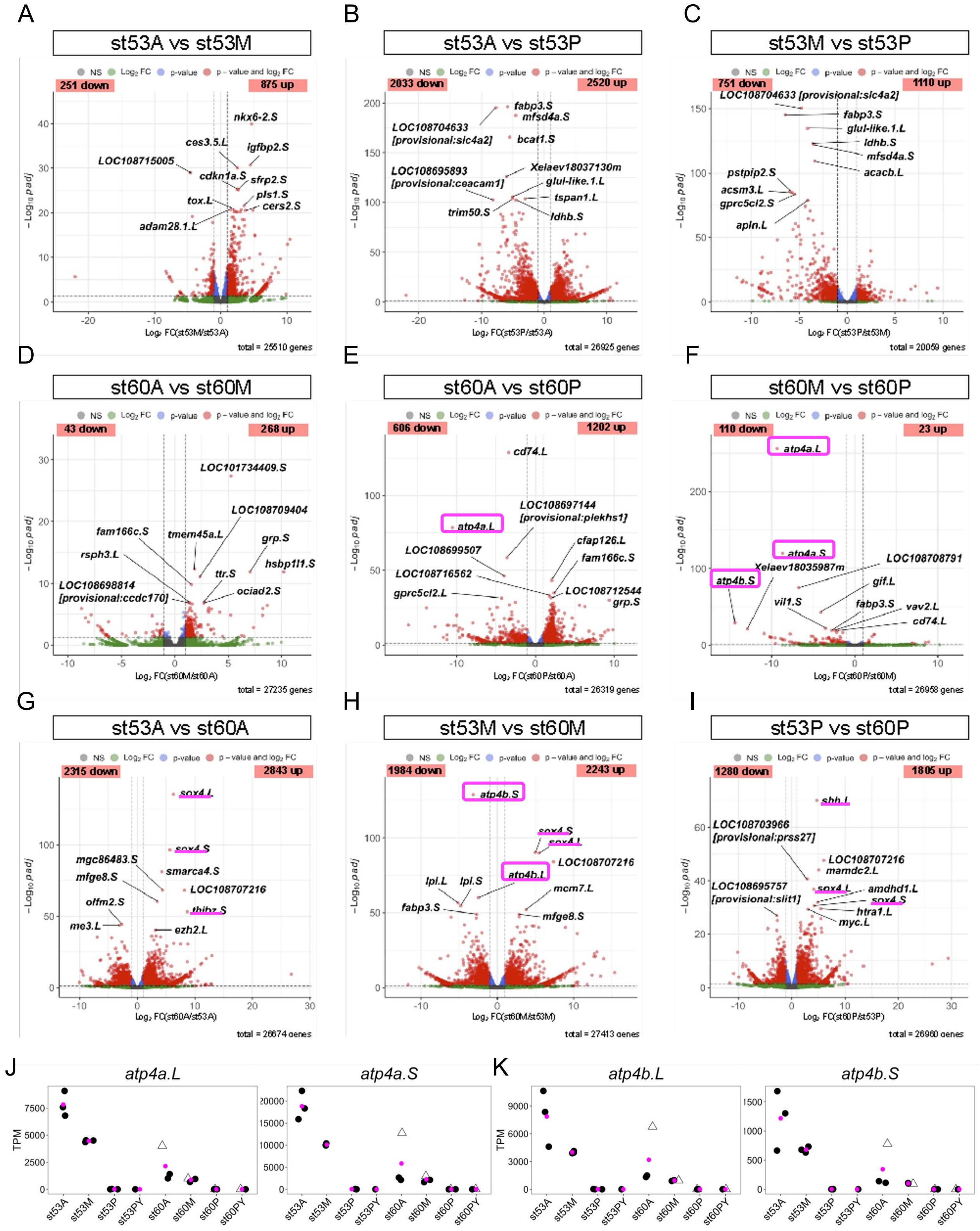

**Fig. S6. Differentially expressed gene analysis of RNA-seq data between pre-metamorphic and metamorphic climax stages.**

(A-I) Volcano plots of differentially expressed genes (DEGs) between anterior (A), mid (M), and posterior (P) regions of the stomach at pre-metamorphic (stage 53) and metamorphic climax (stage 60) stages are shown. The numbers of up- and down-regulated DEGs (e.g. A vs M means up- and down-regulation in M compared to A) between two regions or stages are indicated by the numbers with 'up' or 'down' in the plots. X-axis, log2 fold change; y-axis, -log10 (padj). Red dots, DEGs. The top 10 DEGs from the highest -log10 (padj) value are labeled with gene names. (A-F) Comparison between regions A, M, and P at stage 53 (A-C) and at stage 60 (D-F). Note that the numbers of up- and down-regulated DEGs are both much smaller at stage 60 than at stage 53 (Compare panels A and D, panels B and E, and panels C and F; see also Table S2). The total number of genes plotted are indicated in each panel. This may reflect that regional differences are equalized by dedifferentiation and remodeling of the entire stomach during metamorphosis (see the text). (G-I) Comparison of regions A, M, or P between stages 53 and 60. The numbers of both up- and down-regulated DEGs of stages 53 vs 60 in each region are larger than those between regions at stage 53 or 60 (A-F). Noticeable DEGs at stage 60 are *sox4.L/S*, *thibz.S*, and *shh.L* as up-regulated genes (underlined in G,H,I) and *atp4b.L/S* as down-regulated genes (magenta box in E,F,H). *Sox4.L/S, thibz.S,* and *shh.L/S* are reportedly upregulated by thyroid hormone (TH) in the entire stomach at stage 60 (see also Figs. S8M,N, S9A). (J, K) Dot plots of *atp4a.L/S* (J) and *atp4b.L/S* (K). *atp4a* and *atp4b* encode gastric H(+)-K(+)-ATPase subunits alpha and beta, respectively, which form a proton pump to generate acidic gastric juice in the stomach body; therefore, *atp4a* and *atp4b* genes are used as terminal differentiation markers for the stomach body (Willet and Mills, 2016). Down-regulation of *atp4a.L/S* and *atp4b.L/S* indicates the elimination of differentiated cells and/or the transition to a dedifferentiation state and is an example of the equalization of gene expression profiles in the stomach during metamorphosis. Y-axis, transcripts per million (TPM).

### Fig. S7 (Nagura et al.)

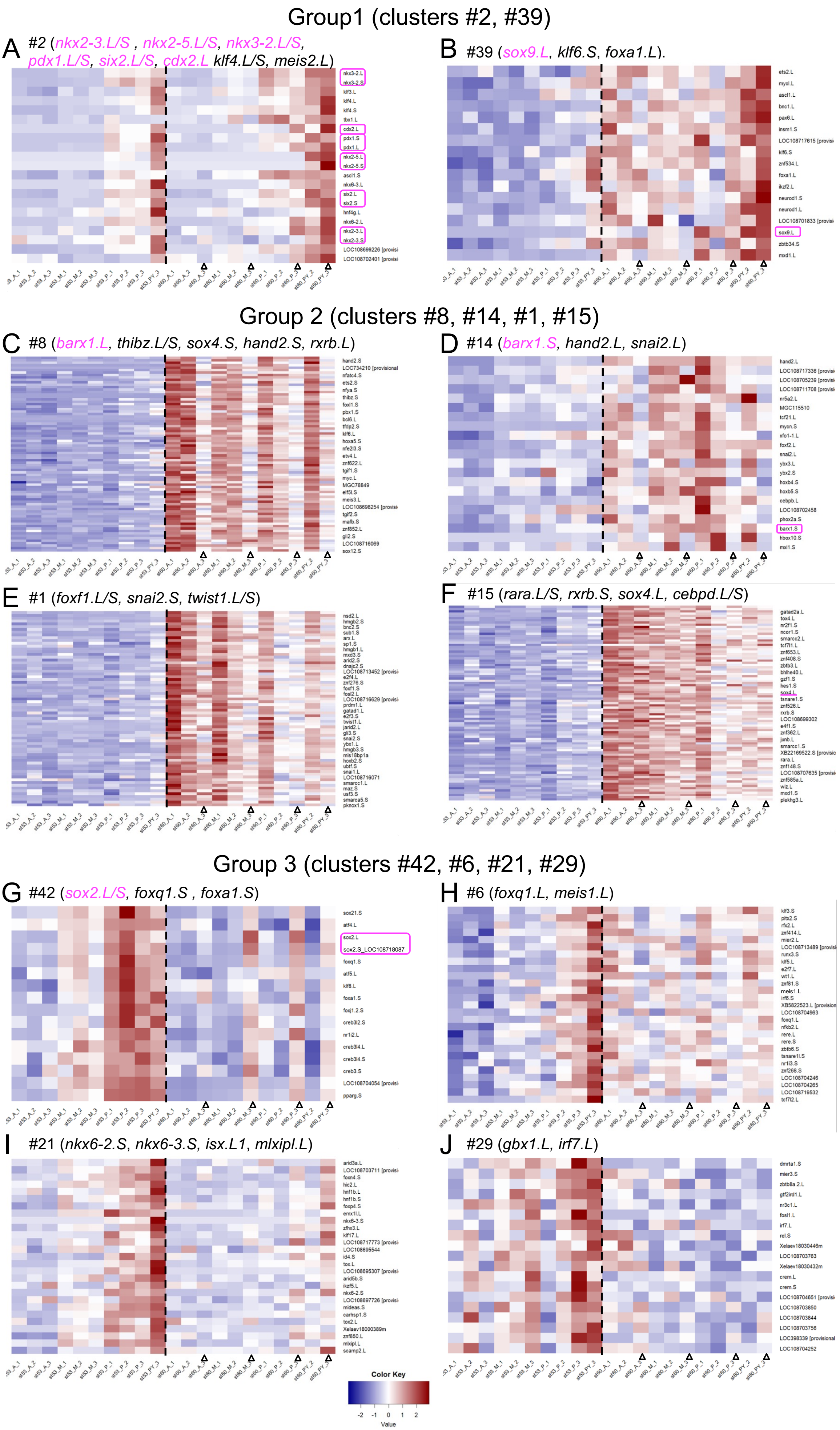

**Fig. S7. Selected clusters of transcription factor genes identified by hierarchical clustering analysis of regional expression profiles in the stomach at pre-metamorphic and metamorphic climax stages.**

Transcription factors (TF) genes with 2 or more TPM (1,252 out of 2,435 genes) were divided into 50 clusters by hierarchical clustering analysis (HCA) (see Fig. 8E; Table S5), and 10 out of them were assembled into three groups (Tables S5, S6; see the main text for details). (A,B) Group 1 gene clusters, #2 and #39. (C-F) Group 2 gene clusters, #8, #14, #1, and #15. (G-J) Group 3 gene clusters, #42, #6, #21, and #29. Representative genes are shown in the parentheses. The pylorus/sphincter-related genes are indicated in magenta and by magenta boxes. Note that too many genes are not all listed on the right side of the heatmaps (C,E,F). See Table S5 for all clusters and their consisting genes. Expression level is shown by color-coded z-scored TPM. Vertical dotted line separates pre-metamorphic stage 53 and metamorphic climax stage 60 samples.

Fig. S8 (Nagura et al.)

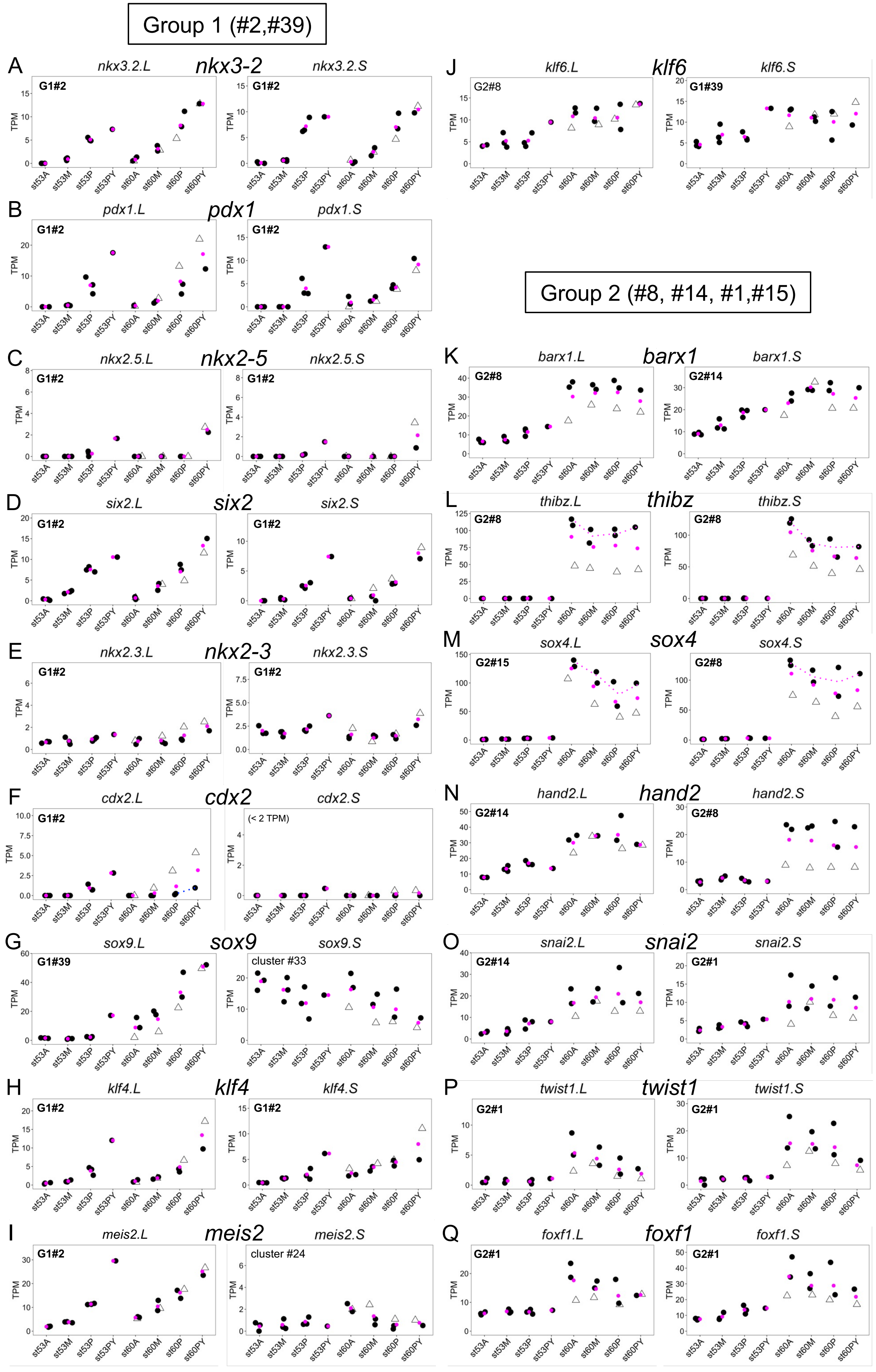

### Fig. S8 (Nagura et al.), continued

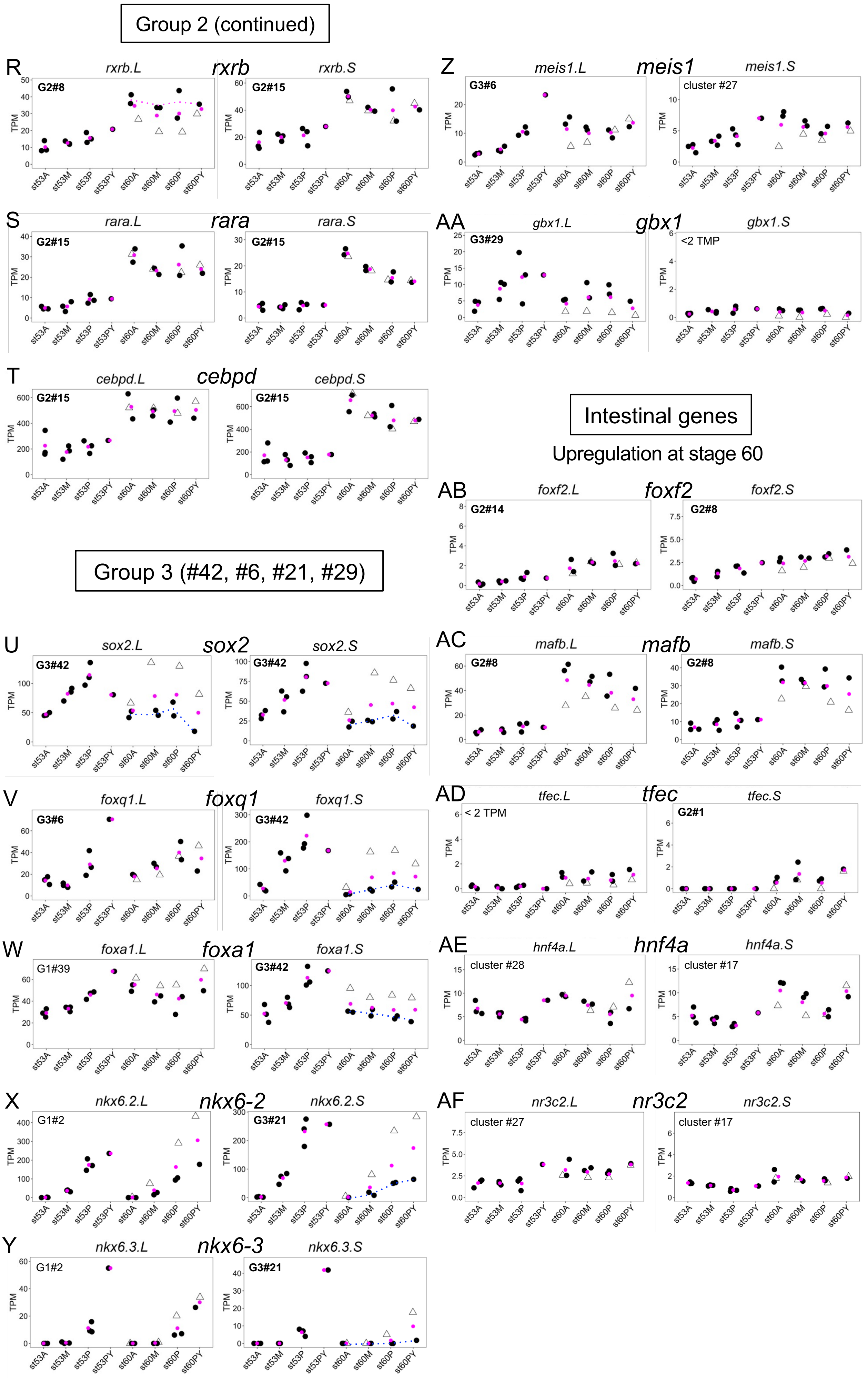

Fig. S8 (Nagura et al.), continued

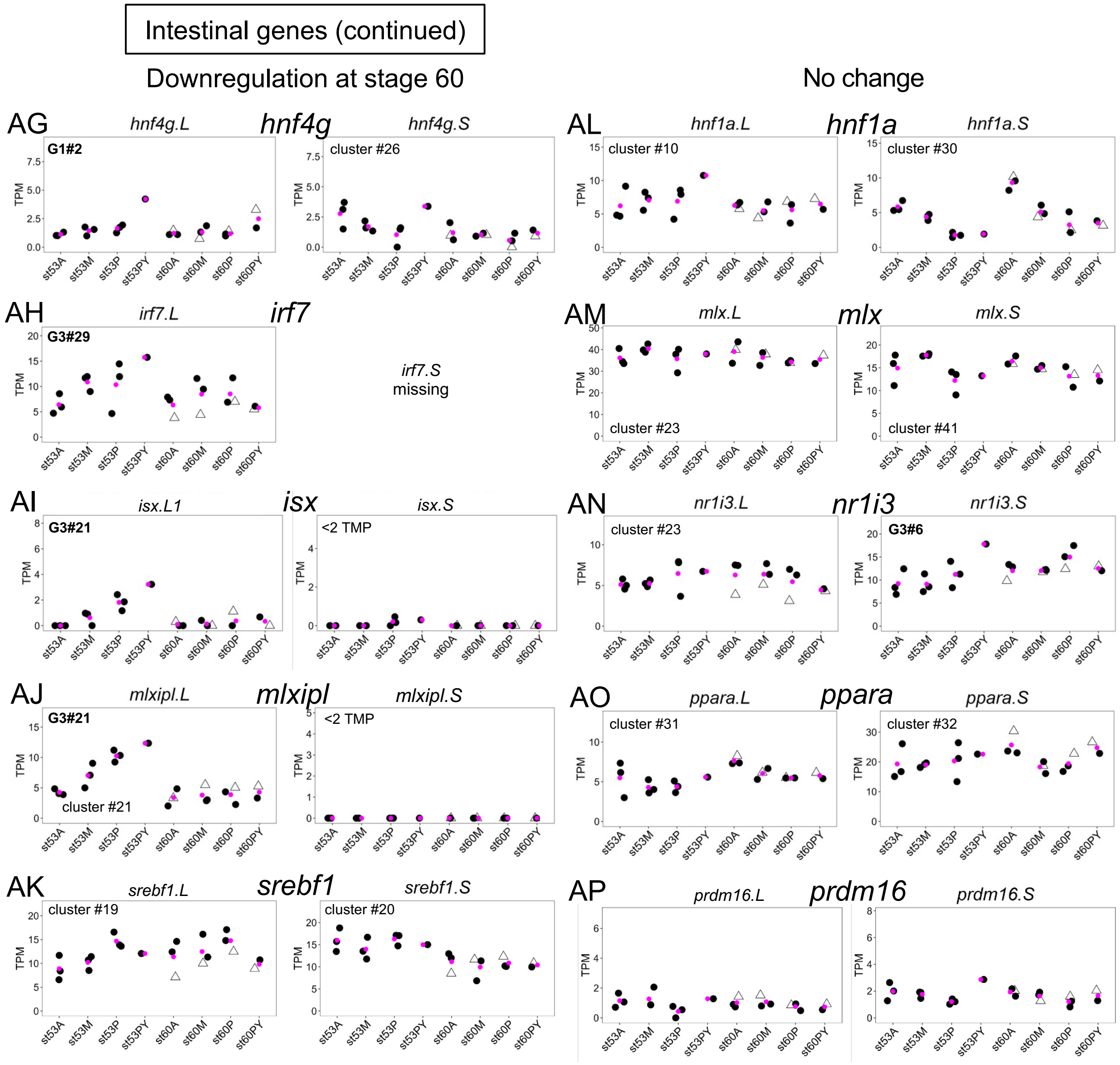

**Fig. S8. Dot plot analysis of regional and temporal expression profiles of transcription factor genes in the stomach at pre-metamorphic and metamorphic stages.**

Fig. S9 (Nagura et al.)

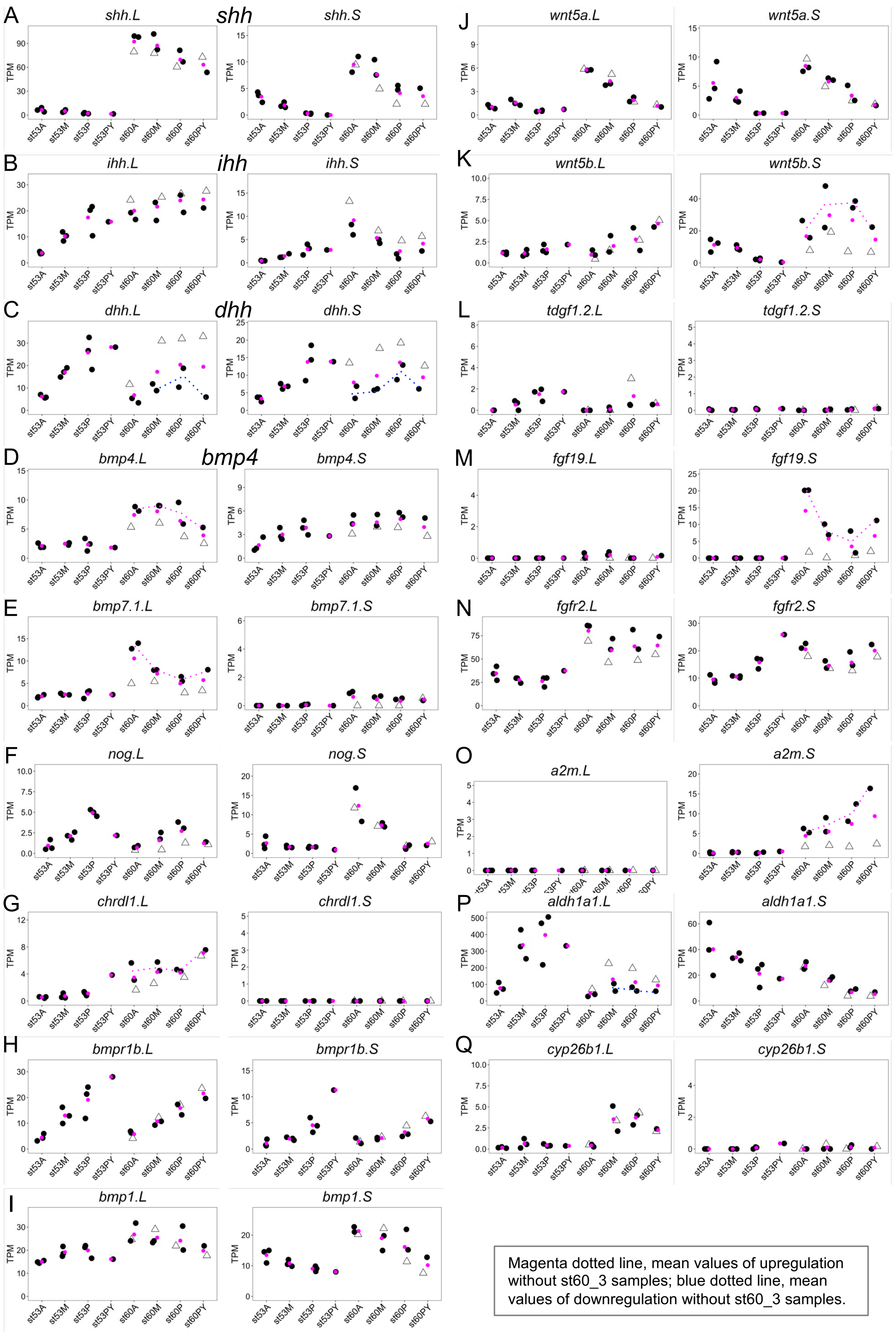

Fig. S9. Dot plot analysis of regional and temporal expression profiles of signaling factor genes in the stomach at pre-metamorphic and metamorphic stages.

Dot plots of expression profiles are shown for selected signaling factor genes. The ordinate, transcripts per million (TPM). The sample names, st53A, st53M, st53P, st53PY, st60A, st60M, st60P, st60PY are indicated below the plot. (A-C) Hh, (D-I) BMP, (J-L) Wnt, (M-O) FGF, (P, Q) Retinoic acid (RA) related genes. (A) *shh*, (B) *ihh*, (C) *dhh*, (D) *bmp4*, (E) *bmp7-1*, (F) *nog*, (G) *chrdl1*, (H) *bmpr1b*, (I) *bmp1*, (J) *wnt5a*, (K) *wnt5b*, (L) *tdgf1-2*, (M) *fgf19*, (N) *fgfr2*, (O) *a2m* (*alpha-2-macroglobulin*; the same as *endodermin*), (P) *aldh1a1*, (Q) *cyp26b1*. All ligand genes for Hedgehog, TGF-β superfamily, Wnt, and FGF signaling that are expressed at a maximum of 2 TPM or higher are shown (A-E, J, K, M). Left and right plots in each panel are L and S homeologs in *Xenopus laevis*. Mean TPM values are shown with magenta dots. Open triangles, st60_3 samples; dotted lines in some panels, average values of upregulation (magenta) or downregulation (blue) without st60_3 samples. For example, *bmp4.L, bmp7.L, chrdl1.L, wnt5b.S, fgf19.S, a2m.S* (D, E, K, M, O), st60_sample 3 (open triangle) showed lower values than st60_samples 1 and 2 at all positions, suggesting that their upregulation occurs at late stage 60. By contrast, *shh.L* and ihh.L (A,B), for example, st60_sample 3 showed values similar to st60_samples 1 and 2, suggesting that their upregulation occurs at early stage 60.
